# Neuronal processes contain the essential components for the late steps of ribosome biogenesis

**DOI:** 10.1101/2025.04.28.650935

**Authors:** Claudia M. Fusco, Anja Staab, Ashley M. Bourke, Georgi Tushev, Kristina Desch, Erico Moreto Lins, Elena Ciirdaeva, Susanne tom Dieck, Nina Kaltenschnee, Alexander Heckel, Julian D. Langer, Erin M. Schuman

**Affiliations:** Max Planck Institute for Brain Research, Max von Laue Strasse 4, 60438 Frankfurt, Germany; Institute for Organic Chemistry and Chemical Biology Goethe-University Frankfurt Max-von-Laue Str. 9, 60438 Frankfurt, Germany; Present address: Developmental Biology Unit, EMBL, Meyerhofstraße 1, 69117 Heidelberg, Germany

**Keywords:** ribosome, protein synthesis, ribosome biogenesis, neurons, local translation

## Abstract

Neurons rely on spatial and temporal control of protein synthesis to respond rapidly and locally to external stimuli, a process facilitated by the dynamic localization and modification of ribosomes. While previous research has shown that neuronal activity can regulate ribosome localization and modify translation rates, little is known about ribosomal assembly within neuronal processes. Here, we investigated the potential for local ribosome maturation in rat neurons using proteomics, RNA sequencing, and imaging methods. We detected an abundance of ribosome biogenesis factors (RBFs) in distal neuronal compartments, particularly those associated with the late stages of ribosome assembly. Moreover, we detected cytosolic pre-rRNA species in dendrites, alongside the enzymes necessary for their processing, suggesting that local ribosome maturation can occur far from the nucleus. These findings challenge conventional models that confine ribosome biogenesis to nuclear and perinuclear regions and suggest that neurons may fine-tune local protein synthesis by regulating ribosome assembly near synaptic sites. This mechanism may enable rapid modulation of the translational capacity in response to physiological changes, regulating synaptic plasticity and local protein synthesis in neurons.

**Significance Statement:** Neurons require precise spatial and temporal regulation of protein synthesis to adapt rapidly to external stimuli, particularly at synapses. Our study challenges the view that new ribosomes can be made exclusively near the nucleus and reveals that ribosome biogenesis factors and pre-rRNA processing enzymes are present in distal neuronal compartments. These findings suggest that ribosome maturation can occur locally in dendrites, enabling rapid, spatially targeted modulation of translational capacity. This mechanism provides a different framework for understanding how neurons regulate synaptic plasticity and adapt to physiological changes. By demonstrating that ribosome assembly may extend beyond the nucleus, this work highlights a previously unrecognized layer of neuronal protein synthesis control, potentially transforming our understanding of how neurons orchestrate local responses to environmental cues.

## Introduction

All cells adapt to their environment through dynamic adjustments to their proteome. Neurons, with their extensive spatial compartmentalization and long lifespan, use a variety of mechanisms to regulate the spatial and temporal availability of proteins. A critical aspect of this regulation is the distribution of the translational machinery (ribosomes) within neuronal processes (1–5), linking protein synthesis directly to neuronal activity and ensuring rapid, localized responses to external stimuli (6, 7).

Neurons also fine-tune local protein synthesis by regulating the local mRNA pool and modifying the activity of translation factors (8–11). Importantly, the ribosome itself can also be a target of regulation (12). For example, neuronal activity leads to the phosphorylation of Ribosomal Protein S6 (RPS6), affecting the translation of specific mRNAs (13, 14). Additionally, increased production of ribosomal proteins (RPs) and the resulting altered ribosome composition leads to the impaired translational homeostasis observed in neurodevelopmental disorders, like the Fragile X syndrome (15). Furthermore, neurons can enhance their local translation capacity in response to activity, by regulating ribosome trafficking (16) and changing their localization - for example - from dendritic shafts to spines or from granular to free states (2, 17). Relatively little is known, however, about whether neuronal processes possess the capacity for core compositional rearrangements of local ribosomes.

The mRNAs encoding ribosomal proteins (RPs) have also been observed in neuronal processes (18–24). We and others have shown that RPs synthesized within neurites (dendrites and axons) can associate with local, pre-existing ribosomes (25, 26). This dynamic RP incorporation could be used for ribosome repair and/or specialization. In fact, the RP incorporation probability depends on the subcellular location and physiological state. For example, the incorporation of a number of nascent RPs was increased after oxidative stress (25) or after stimulation with a growth factor (26). Similar findings have recently been observed in yeast, where one of the exchanging RPs (RPS26) gets preferentially damaged under oxidative stress, and replaced by a new functional copy (27, 28). During this process, ribosomes that transiently lack RPS26 acquire a new translational preference for genes related to the stress-response (29). Neurons may use similar mechanisms to dynamically modify the translational machinery near synapses. Additionally, there is the possibility that distally synthesized RPs can be incorporated into premature ribosomes, facilitating their maturation and thereby expanding the local pool of functional ribosomes. This idea of local ribosome maturation, however, challenges the prevailing view that immature ribosomal particles are only found in nuclear and perinuclear regions.

Ribosome biogenesis begins with ribosomal RNA (rRNA) transcription in the nucleolus and involves the coordinated association of several ribosome biogenesis factors (RBFs), RPs, and small nucleolar RNAs (snoRNAs). Such interactions allow pre-rRNA processing (cleavage), modification (pseudouridylation and methylation) and folding into pre-ribosomal subunits (30). This process continues in the nucleoplasm and ends in the cytoplasm, where the small (40S) and large (60S) ribosomal subunits become competent for mRNA translation (31). Although the overall process is highly conserved, the discovery of unique regulatory features in mammalian cells has highlighted a more complex assembly pathway in higher eukaryotes (32). Most RBFs transiently or stably associate with forming pre-ribosomal particles and trigger conformational rearrangements at specific steps either in the nucleolus, nucleoplasm or cytoplasm. Maturation is then completed when all RBFs are removed and all RPs are incorporated (32). Throughout the process, several quality control mechanisms ensure the proper folding and functional competence of mature ribosomes (33). However, whether ribosome biogenesis, particularly its later stages, is modulated by physiological conditions, remains poorly understood (34). Instead, most research has focused on the regulation of the early steps, such as rRNA transcription and RP translation (15), often in relation to cellular proliferation and cancer through pathways like mTOR signaling (35, 36).

Here we used imaging, mass spectrometry, and RNA sequencing to characterize markers of ribosome biogenesis in different neuronal subcellular compartments. We discovered that particles from the late (but not early) stages of ribosome biogenesis are localized in distal neuronal processes. Our findings suggest a novel mechanism by which neurons may modify their translation capacity through the local maturation of ribosomes.

## Results

### The subcellular distribution of translation-related proteins and ribosome biogenesis factors

To characterize, in an unbiased manner, the distribution of translation-related proteins between subcellular compartments, we cultured rat cortical neurons in compartmentalized chambers that allow for the enrichment of a neurite (axon + dendrite) fraction for comparison with a mixed population of somata and neurites (Figure 1a). We then separately purified proteins from either compartment and used Data Independent Acquisition (DIA) Mass Spectrometry to characterize the subcellular proteomes (Figure 1b). We reliably identified ∼9000 and ∼8000 protein groups in the somata+neurite and neurite-only samples, respectively (Suppl. Figure 1). We compared the proteome from these two compartments and found 2594 proteins enriched in the somata+neurite fraction and 944 proteins enriched in the neurite-only fraction (Figure 1c). Gene ontology analysis of the differentially regulated proteins confirmed the specificity of our preparation: terms associated with the nucleus or the perinuclear region (e.g., the spliceosome and Endoplasmic Reticulum, respectively) were enriched in the somata+neurite fraction, while synaptic terms were enriched in the neurite-only fraction (Figure 1d). To examine how neurons distribute their translational machinery between subcellular compartments, we curated a list of gene categories related to protein synthesis including ribosomal proteins, initiation factors, elongation factors, RNA binding proteins, and ribosome biogenesis factors. We determined the degree to which a given category was enriched in either compartment, relative to the other. When analyzed in this way ribosomal proteins showed a strong de-enrichment in neurites and a strong enrichment in somata+neurites (Figure 1e and d). Interestingly, initiation and elongation factors were, in contrast, equally distributed between compartments (Figure 1e). These two observations are consistent with the recent observation that protein synthesis in the cell body is mostly undertaken by polysomes (several ribosomes translating the same mRNA), while neuronal processes tend to prefer protein synthesis by monosomes (one single ribosome per mRNA) (19). The higher ratio of translation factors per ribosome in the neurite compartment fits with the slower initiation and elongation rates exhibited by monosome-preferring transcripts, as they require more time to process a higher number of secondary structures in the mRNA (19). This is paralleled by the apparent higher proportion of RNA binding proteins per ribosome in the neurite compartment (Figure 1e). Altogether, our data suggests that neuronal processes have a high capacity to regulate RNA localization and translation by differentially distributing key regulators of protein synthesis.

**Figure 1.**
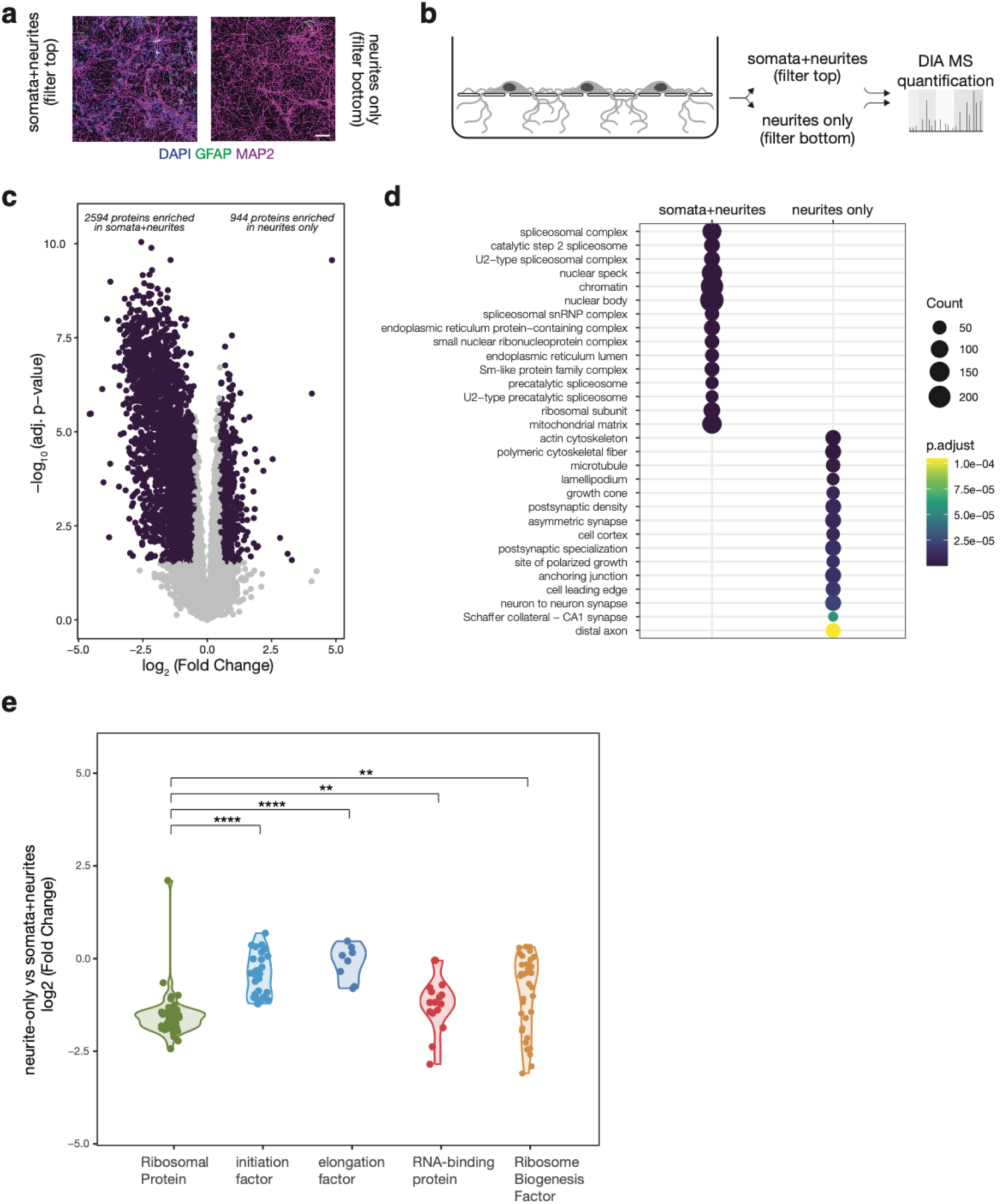
Local proteome profiling reveals the subcellular distribution of translation-related proteins. (**a**) Representative images indicating the presence of dendrites (anti-MAP2; top and bottom), nuclei (DAPI; top only) and glia (anti-GFAP; top only) in the top or bottom sections of a compartmentalized chamber, as indicated (scale bar = 50 μm). (**b**) Schematic of the experimental design for measuring protein abundance between subcellular compartments. The top and bottom of each filter was separately harvested for downstream data-independent acquisition (DIA) mass spectrometry (MS). (**c**) Volcano plot of significantly regulated proteins (purple, FDR<0.05 and >|0.5| log-scaled fold change) between total lysates from the somata+neurite and neurites-only fractions. Unpaired, two-sided *t*-test with Welch correction on rows. Benjamini-Hochberg correction for multiple testing. (**d**) Gene Ontology (GO) analysis of Cellular Components terms overrepresented (FDR<0.01) among the differentially regulated proteins between total lysates from soma+neurites and neurites-only fractions. (**e**) Violin plot of the log2 fold change of the protein abundance between the neurite-only and soma+neurite fractions, grouped by gene category. Each dot represents a protein of interest. The number of detected proteins in each group was 47, 29, 8, 16, and 36 for ribosomal proteins (RP), initiation factors, elongation factors, RNA-binding proteins (RBP) and Ribosome biogenesis factors (RBF), respectively. Kruskal−Wallis, p < 2.2e−16; *t*-test between RP and initiation, p = 1.8e−14; *t*-test between RP and elongation, p = 2.2e−07; *t*-test between RP and RBP, p = 0.004; *t*-test between RP and RBF, p = 0.0017.

The above proteomic characterization also revealed a surprisingly high proportion of RBFs in the neurite compartment (Figure 1e). To explore this further, we asked whether the detection and abundance of a RBF in the neurite compartment is influenced by the location (nuclear vs. cytoplasmic) in which it is known to interact with immature ribosomes. Indeed, most RBFs that dissociate from the pre-ribosomal subunits in the nucleus (see Figure 3a) were detected only in the somata+neurite compartment. In contrast, almost all the RBFs that dissociate from the pre-subunits in the cytoplasm were detected in both compartments (Figure 2a). Importantly, when detected in the neurites, nuclear RBFs displayed lower levels than their cytoplasmic counterparts (Figure 2b-c and Suppl. Fig.1d-f).

**Figure 2.**
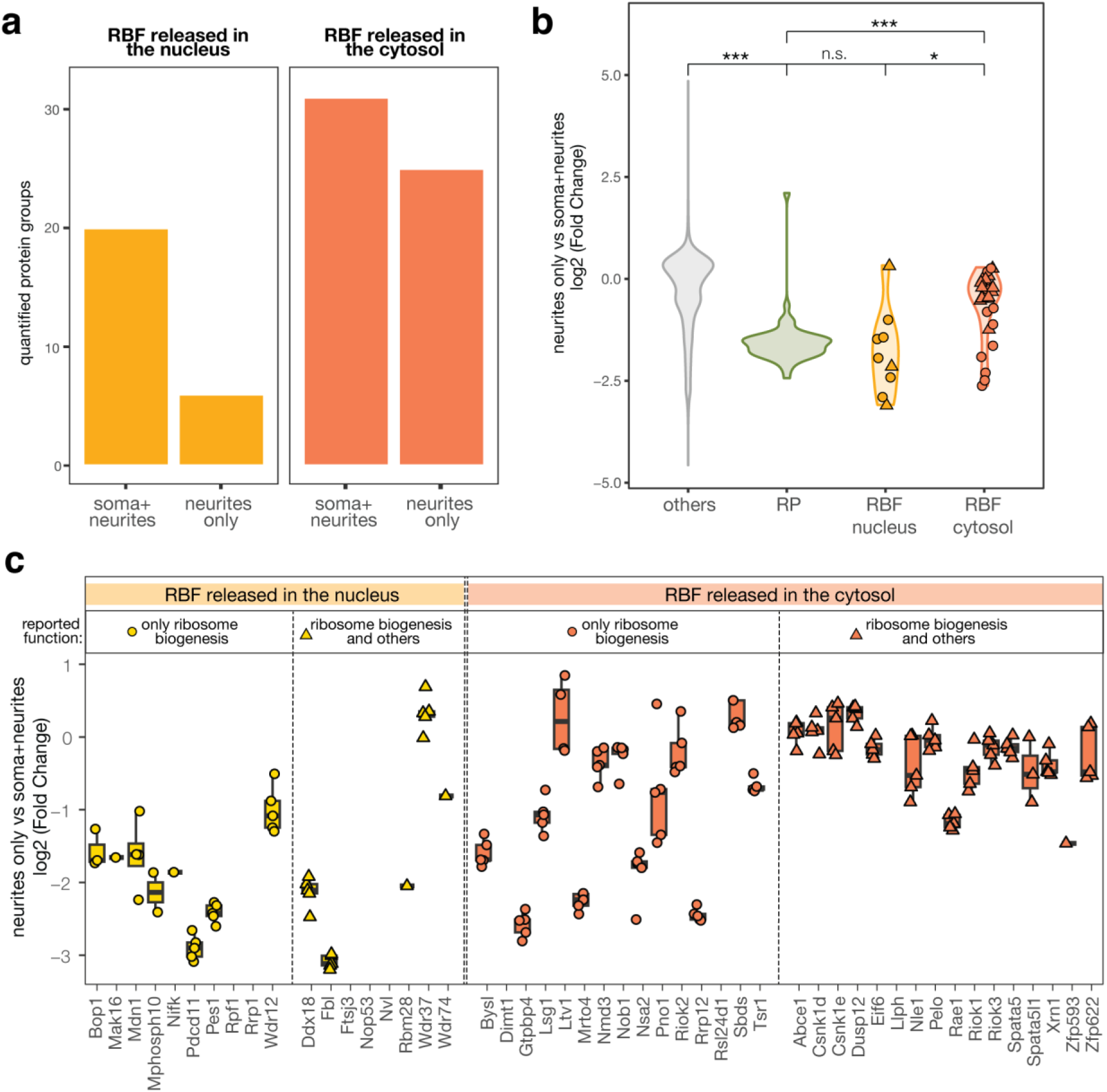
Cytosolic (but not nuclear) ribosome biogenesis factors localize to neuronal processes. (**a**) Bar plot of the number of identified RBFs in all biological replicates (5/5) of each subcellular compartment. The bars related to RBFs that are released in the nucleus are colored light orange, those for RBFs that are released in the cytoplasm in dark orange. (**b**) Violin plot of the log2 fold change of the protein abundance between neurite-only and somata+neurite fractions, as in (a). Each dot represents a RBF. The shape of the dot highlights if the corresponding protein is known to have other (e.g., moon-lighting) functions. RBFs that dissociate from the pre-ribosome subunits in the cytosol (in orange, n = 27), but not those that are released in the nucleus (in yellow, n = 9), show a significantly higher proportion in the neurite-only fraction compared to Ribosomal Proteins (in green, n = 47). Kruskal−Wallis, p < 2.2e−16; *t*-test between RP and others, p < 2.22e−16; *t*-test between RP and RBF nucleus, p = 0.32; *t*-test between RP and RBF cytosol, p = 2.6e−06; *t*-test between RBF nucleus and RBF cytosol, p = 0.0098. (**c**) Boxplot showing the log2 intensity of quantified Ribosome Biogenesis Factors (RBFs) in the neurites only, compared to the soma+neurite compartment. Each point represents a biological replicate. Proteins quantified in one compartment only are shown as missing values. RBFs that are released in the nucleus are yellow, those for RBFs that are released in the cytoplasm in orange. The shape of the dot indicates if the corresponding protein is known to have extra-ribosomal (e.g., moonlighting) functions. Abbreviations: Ribosomal Proteins (RP), RNA Binding Protein (RBP), Ribosome Biogenesis Factors (RBF), repl. (biological replicate).

**Figure 3.**
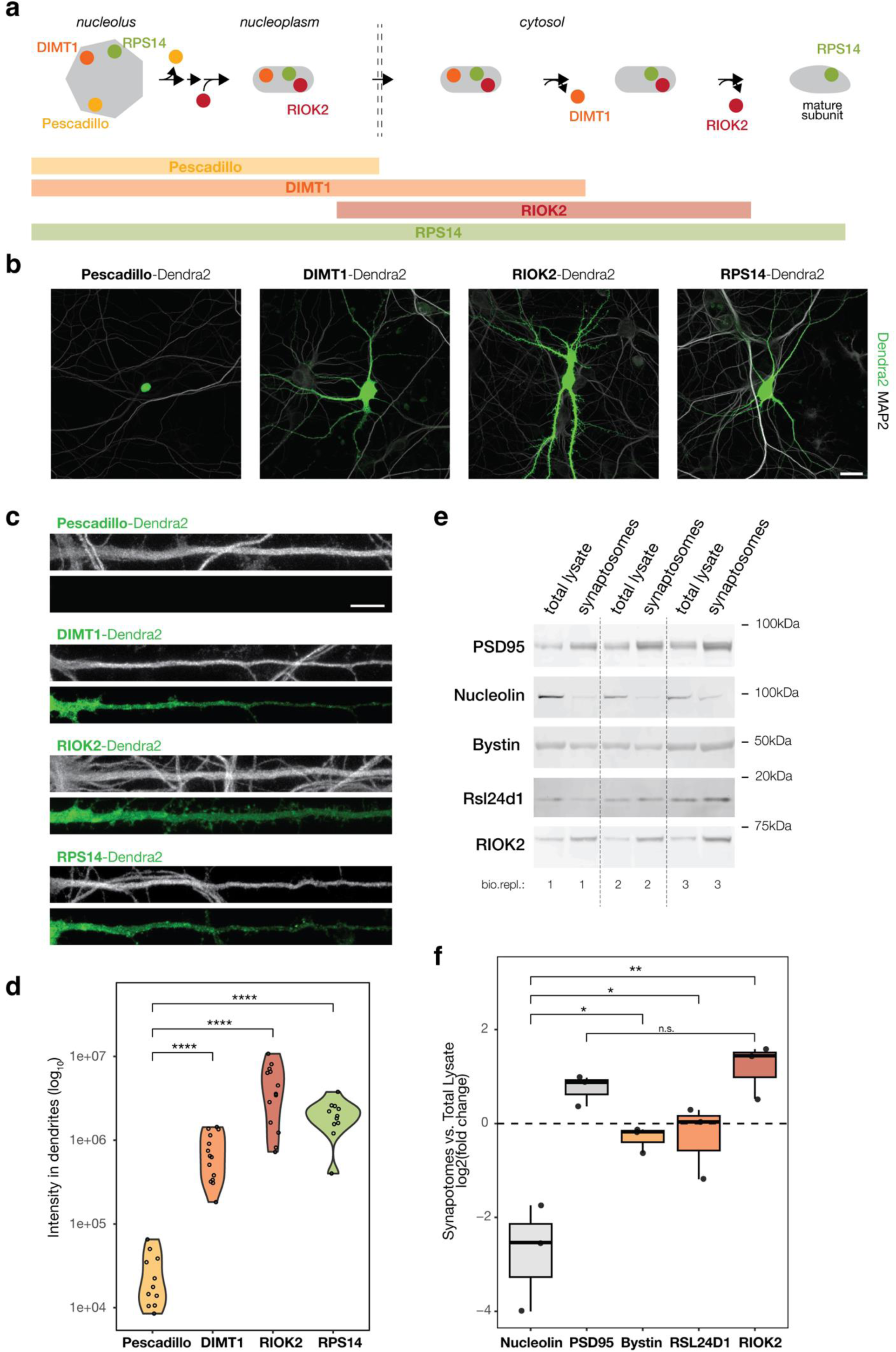
Cytosolic (but not nuclear) ribosome biogenesis factors localize to neuronal processes and synapses. (**a**) Schematic representation of the incorporation and dissociation of different biogenesis factors (Pescadillo, DIMT1, RIOK2, respectively in yellow, orange and red) and a Ribosomal Protein (RPS14, in green) in different cellular compartments (from left to right: nucleolus, nucleoplasm and cytosol) during the maturation of the ribosomal subunits (in gray). (**b**) Representative images showing the localization (within individual neurons) of different biogenesis factors (Pescadillo, DIMT1, RIOK2) and a Ribosomal Protein (RPS14) fused with the common fluorescent reporter Dendra2 (scale bar = 25μm). The dendritic marker MAP2 is shown in grey, Dendra2 in green. (**c**) Straighten dendrites of neurons shown in Fig. 3b (scale bar = 10μm). All dendrites are shown with the same orientation (part most proximal to the soma is on the left, distal to the right). The dendritic marker MAP2 is shown in grey, Dendra2 in green. (**d**) Violin plot depicting the Dendra2 intensity of the indicated proteins in the dendritic arbor of individual neurons. n= number of cells, was between 11 and 14 (as indicated by the dot plots), over 3 independent experiments. Kruskal−Wallis, p = 4.3e−08; *t*-test between Pescadillo and DIMT1, p = 4.5e−07; *t*-test between Pescadillo and RIOK2, p = 8e−07; *t*-test between Pescadillo and RPS14, p = 2.8e−06. (**e**) Detection of proteins-of-interest in total lysate and synaptosome fractions from rat cortex by Western blot (three biological replicates). (**f**) Quantification of Western blot signals, showing the log₂ fold change in normalized protein levels (relative to total protein staining) between synaptosome and total lysate fractions. The nuclear marker Nucleolin was strongly depleted in synaptosomes, while the synaptic marker PSD95 was enriched, confirming the purity of the preparation. All three tested RBFs—RSL24D1, Bystin, and RIOK2—were present in synaptosomes at levels exceeding those of Nucleolin (ANOVA, p = 0.00038; paired t-test: ENP1, p = 0.043; RSL24D1, p = 0.019; RIOK2, p = 0.0077). The enrichment of RIOK2 in synaptosomes was statistically indistinguishable from that of PSD95 (paired t-test, p = 0.43).

To validate the unexpected detection of RBFs in neuronal processes, we created fluorescent protein fusion constructs and assessed their subcellular localization following expression in neurons. We selected three RBFs and one RP that participate in the maturation of ribosomal subunits at well-characterized check-points and for which no other function besides ribosome biogenesis has been reported (Figure 3a). As a “nuclear only” control, we chose the protein Pescadillo which is known to incorporate into pre-60S particles in the nucleolus and dissociate once in the nucleoplasm (37). As cytoplasmic RBFs we chose DIMT1 and RIOK2. DIMT1 associates with the small subunit processome in the nucleolus and is released in the cytoplasm (38, 39). RIOK2 is incorporated into the pre-small subunit just prior to nuclear export and is one of the last factors to be released in the cytoplasm as biogenesis is completed (32). Finally, as a constitutive element of ribosomes, we chose Ribosomal Protein RPS14, which stably associates with the ribosome early (in the nucleolus) and remains a core element of the small ribosomal subunit for its lifespan (40). To avoid difficulties that might arise from differences in antibody-epitope binding, we fused each protein-of-interest to the fluorescent reporter Dendra2, and transfected cultured neurons. One day after transfection, we imaged individual neurons and quantified the distribution of each protein-of-interest in the nucleus, and in the somatic and dendritic cytoplasm. As expected, Pescadillo was detected only in the nucleus (both nucleolus and nucleoplasm), while DIMT1, RIOK2 and RPS14 were also detected in the cytoplasm (Figure 3b). Importantly, the cytoplasmic signal of both DIMT1 and RIOK2 was not confined to the soma but distributed throughout the dendritic arbor (Figure 3c-d). The signal for RIOK2 was especially prominent at putative synaptic sites.

To further assess the presence of ribosome biogenesis factors (RBFs) at synapses, we analyzed synaptosome preparations from rat cortex. Western blot analysis confirmed the enrichment and depletion of synaptic and nuclear material, as validated by the presence of the synaptic marker PSD95 and the absence of the nuclear marker Nucleolin, respectively. We then quantified three RBFs—RSL24D1, Bystin, and RIOK2—that associate sequentially with pre-ribosomal particles in the nucleus and are later released into the cytosol (32). Importantly, none of these proteins have reported functions outside of ribosome biogenesis. All three RBFs were detected in the synaptosome fractions at levels exceeding those of nuclear markers, with RIOK2 reaching enrichment levels comparable to PSD95 (Figure 3 e-f).

Altogether, our imaging and biochemical data corroborate the observation that cytosolic and late-stage RBFs populate neuronal processes and synapses.

*RBFs from neurites are assembled in particles*.

The RBFs we observed in dendrites have not been reported to have any other function outside ribosome biogenesis (Figure 2, 3 and Suppl.Fig.1d-f); as such the dendritic localization of these proteins suggest the presence of pre-mature ribosomes in distal processes. To examine this further, we asked whether RBFs in dendrites are associated with assembled ribosomal particles. To this end, we performed a sucrose cushion from either compartment (somata+neurites vs. neurite-only) and measured the proteins present by DIA Mass Spectrometry (Figure 4a). The sucrose cushion conditions were optimized to enrich not only actively translating ribosomes but also pre-ribosomal particles (Suppl. Figure 2; see methods). In the neurite-only compartment, we reliably quantified 47 RPs which shared a homogenous and strong enrichment in the cushion, confirming the successful ribosomal purification. We also quantified 34 RBFs which showed a broader distribution (Figure 4b). We asked whether the localization and strength of interaction between the RBF and the pre-ribosomal particle could explain the breadth of distribution. Indeed, cytosolic, but not nuclear, RBFs and stable, but not transient ribosome interactors, were enriched in the cushion (Figure 4c and d). Altogether, our data show that RBFs in neuronal processes are stable components of assembled particles.

**Figure 4.**
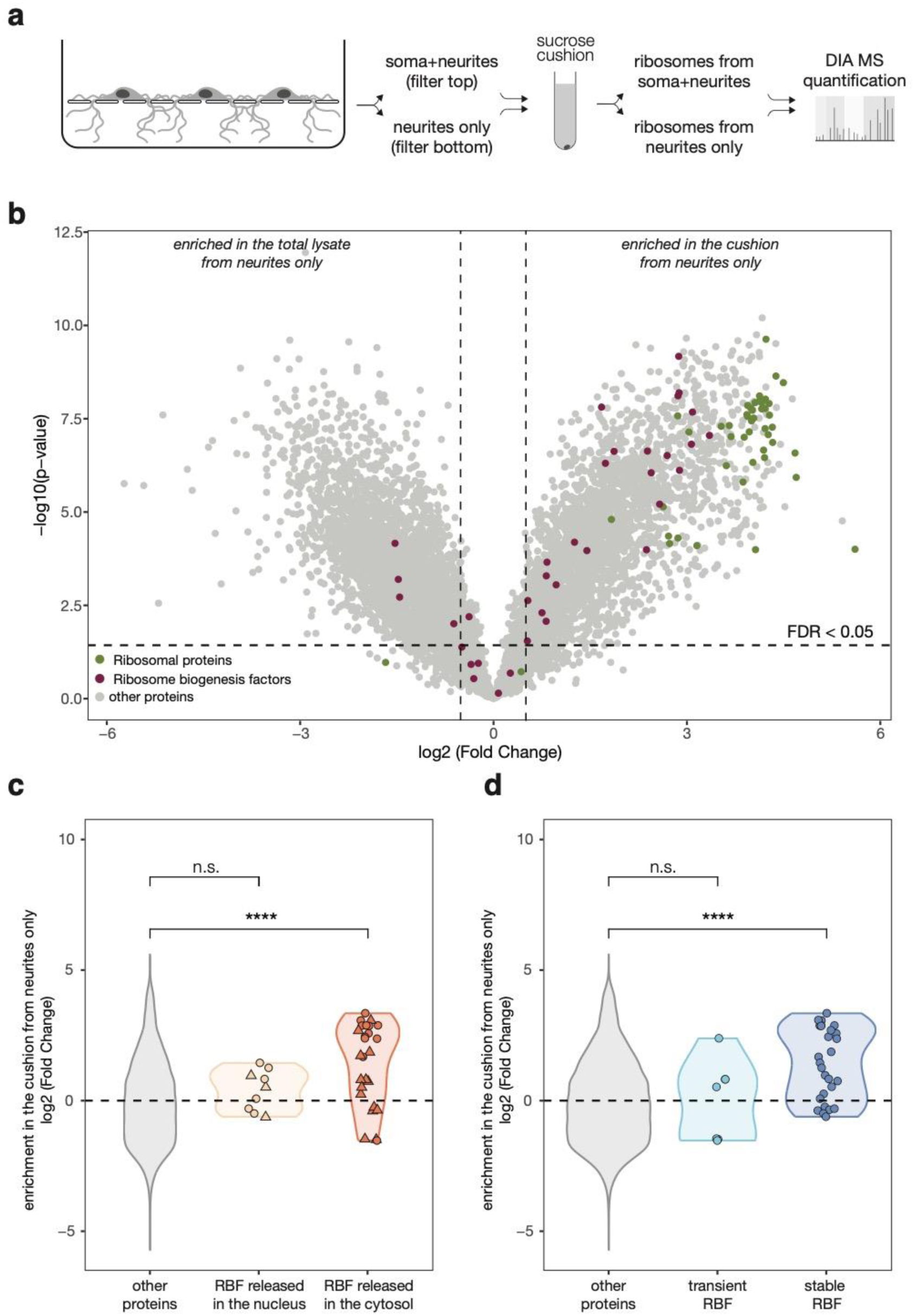
RBFs from neurites are assembled in particles. (**a**) Schematic of the experimental design. Immature and mature ribosomes were purified from either subcellular compartment (somata+neurites vs neurites only) via sucrose cushioning and measured by mass spectrometry. (**b**) Volcano plot of differentially abundant proteins between total lysates and cushion from neurites. Unpaired, two-sided *t*-test with Welch correction on rows. Benjamini-Hochberg correction for multiple testing. N for ribosomal proteins, ribosome biogenesis factors and other proteins is 47, 35, and 7171, respectively. (**c**) Violin plot of the log2 fold change of the protein abundance in the cushion vs the total lysate, grouped by gene category. Each dot represents a protein of interest. Ribosome Biogenesis Factors that dissociate from the pre-subunits in the cytosol (in orange; n = 26), but not those that are released in the nucleus (in yellow; n = 9), were significantly enriched in the cushion from neurites. Kruskal−Wallis, p = 0.00014; *t*-test between others and RBF nucleus, p = 0.29; *t*-test between others and RBF cytosol, p = 4.6e−05. The shape of the dot indicates if the corresponding protein is known to possess (triangle) or lack (circle) functions distinct from roles in ribosome biogenesis. (**d**) Violin plot of the log2 fold change of the protein abundance in the cushion vs the total lysate, grouped by gene category. Each dot represents a protein of interest. Ribosome Biogenesis Factors that stably associate with pre-ribosomal subunits (in dark blue; n = 27), but not those that are just transient interactors (in light blue; n = 7) were significantly enriched in the cushion from neurites. Kruskal−Wallis, p = 7.5e−05; *t*-test between others and RBF nucleus, p = 0.97; *t*-test between others and RBF cytosol, p = 1.3e−05.

### Cytosolic (but not nuclear) pre-rRNAs localize to neuronal processes

To examine whether these RBF-containing particles are indeed pre-ribosomal subunits, we probed the localization of pre-rRNAs. The rDNA gene is initially transcribed as a long transcript (47S rRNA); the activity of exo- and endo-nucleases generates smaller intermediate molecules, until the 5.8S, 18S and 28S rRNAs are fully matured and competent for translation in the cytosol. Studies from yeast and humans have revealed that major steps of ribosome biogenesis and rRNA processing are relatively conserved throughout evolution: the processing of the 28S rRNA occurs only in the nucleus, while the last steps of 18S and 5.8S rRNA maturation are completed in the cytosol (Figure 5a). Despite the astonishingly high degree of conservation between mature rRNAs, the primary sequence around these regions is not conserved, even just within mammalian species (30) (Figure 5b). We note that detecting these cytosolic precursors in neurons poses significant challenges, requiring methods with i) high sensitivity to detect signals in subcellular compartments with low RNA concentrations, ii) a high dynamic range to distinguish them from the highly concentrated signal in the nucleus, and iii) high resolution to differentiate mature and immature rRNA species which differ by only a few dozen nucleotides at their 3’ end. We tested several approaches, including fluorescence in situ hybridization (FISH), molecular beacons, and PCR without consistent success. As indicated below, highly sensitive Northern blotting and RNA sequencing proved successful for detecting cytosolic pre-5.8S rRNA, although detection of cytosolic pre-18S rRNA remained unsuccessful.

**Figure 5.**
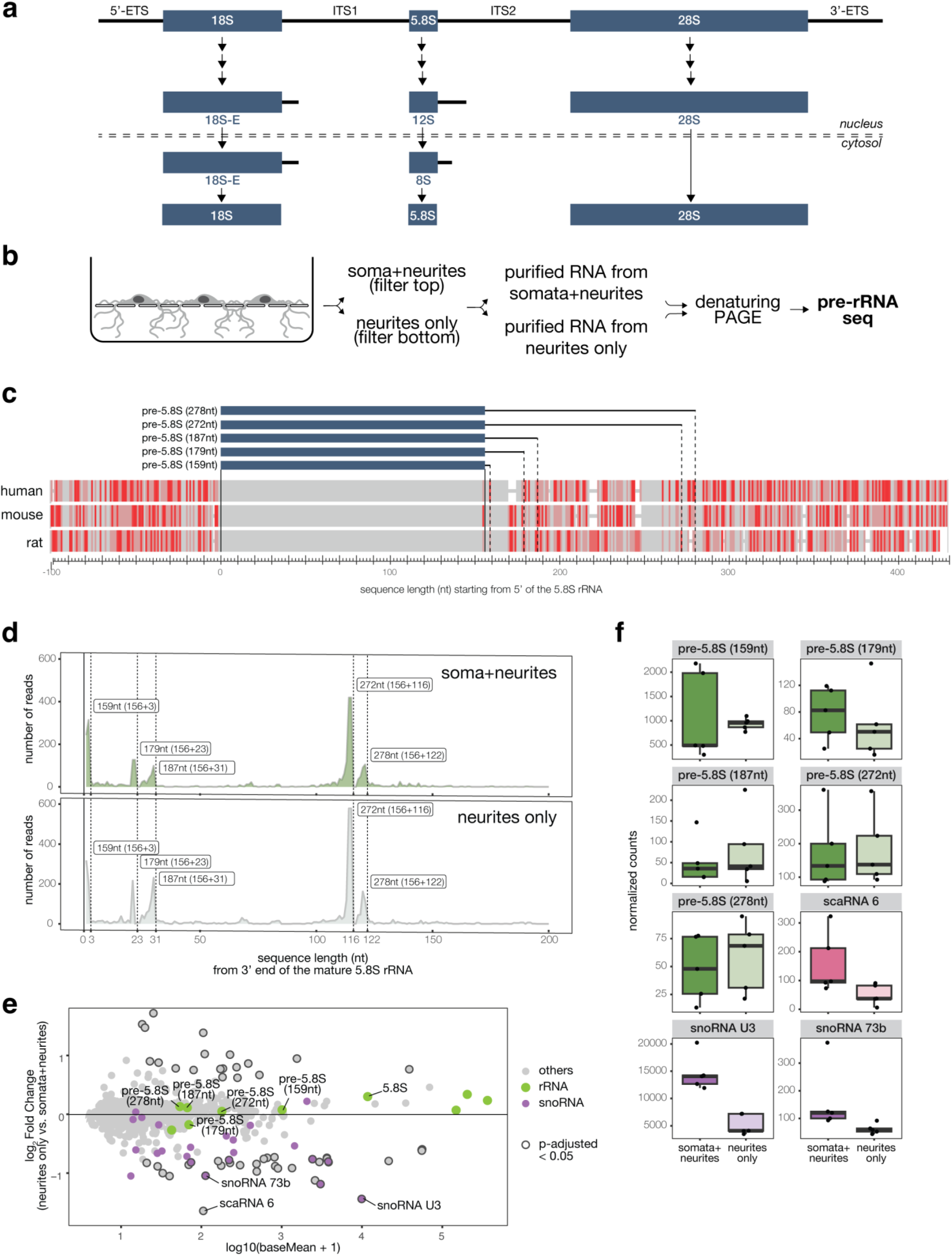
pre-rRNA-seq reveals cytosolic pre-5.8S rRNAs in rat neurons and neurites. (**a**) Schematic representation of the pre-rRNA processing during ribosome biogenesis. Abbreviations: External Transcribed Spacer (ETS), Internal Transcribed Spacer (ITS). (**b**) Multi-sequence alignment of the 100nt upstream and 415nt downstream of the first nucleotide of the mature 5.8S rRNA in human, mouse and rat. Identical nucleotides are shown in gray, mis-matches in red according to the Frequency-base differences method of the NCBI Multiple Sequence Alignment Viewer, Version 1.25.1 (bases that occur more frequently are highlighted in lighter colors). Solid lines indicate the 5’ and 3’ ends of the mature 5.8S. Dashed lines indicate the predicted 3’ end of cytosolic pre-5.8S species identified. (**c**) Schematic of the experimental design. RNAs were purified from either subcellular compartment (somata+neurites vs neurites only) and run on a denaturing TBE-Urea gel to separate RNAs of different sizes. RNA molecules were measured by RNAseq. (**d**) Coverage plot of the 3’ nucleotide of sequenced RNA molecules downstream of the mature 5.8S rRNA. Solid line indicates the 3’ end of the mature 5.8S. Dashed lines indicate predicted 3’ end of the pre-mature 5.8S molecules (estimate of the total length is indicated). On the top (dark green) reads in the somata+neurites sample. On the bottom (light green) reads in the neurite-only sample. Each coverage plot represents the sum of five biological replicates. (**e**) MA plot of the expression levels of RNAs identified in the somata+neurites and neurite-only compartments. Small nucleolar RNAs are shown in magenta, ribosomal RNAs in green. RNAs that are significantly enriched (adjusted p-value < 0.05) in either compartment are highlighted with a dark circle. RNAs of interest are indicated by name. (**f**) Box plot of count of reads mapped to the unique sequence of the cytosolic pre-5.8S molecules, small Cajal-body specific RNA (scaRNA 6), small nucleolar RNA (snoRNA) U3 and 73b, as indicated. Darker colors are used for the somata+neurites samples, lighter colors for the neurite-only samples. Each dot represents a biological replicate.

To identify the sequence of cytosolic pre-5.8S and predict their cleavage sites for the first time in rat tissue, we developed a highly-sensitive pre-rRNA sequencing method (Figure 5c and Suppl. Figure 3a). Total RNA from rat neurons was analyzed using gel electrophoresis to separate RNA molecules according to their size. We then cut gel bands between ∼160 nt and ∼400 nt to enrich for cytosolic pre-5.8S species and de-enrich for mature 5.8S. Purified RNA was then processed to generate roughly 80nt long reads covering the 3’ end of the sequence. As the difference between the mature 5.8S and its cytosolic pre-intermediates is at the 3’ end, the 3’ bias of our sequencing pipeline allowed for the prediction of the exact sequence length and potential cleavage sites of the pre-5.8S species. Additionally, the method was designed to overcome modifications and tertiary structures in the RNA, which are common for rRNA molecules but inhibit most conventional cDNA synthesis pipelines. Consistent with its abundance, the great majority of the reads mapped to the last 80nt of the mature 5.8S rRNA sequence. However, we also quantified a significant number of reads that mapped to the 5’ end of the Internal Transcribed Spacer ITS2 and therefore were unique to pre-5.8S molecules (Figure 5d). In total, we identified five pre-5.8S molecules (159, 179, 187, 272 and 278nt long), three more than what has previously been identified in humans (180nt and 325nt long). Interestingly, when we compared the sequence of the cytosolic pre-5.8S between human, mouse and rat, we observed higher homology at the 5’ of the predicted cleavage sites in rat (Figure 5b), suggesting that despite the overall low similarity in the primary sequence, the general processing and the responsible enzymes might be conserved across mammalian species. We then examined whether these pre-rRNA species can also be found specifically in neuronal processes. To do so, we applied our pre-rRNAseq method to the compartmentalized chambers described above, and identified the above pre-rRNA molecules in the neurite-only compartment, at levels similar to that detected in the somata+neurite compartment (Figure 5e and f). Importantly, nuclear RNAs like the small Cajal body-specific RNA 6 (scaRNA 6) and the small nucleolar RNAs snoRNA U3 and snoRNA 73b, exhibited instead a specific de-enrichment in the neurite compartment, confirming the purity of our preparation (Figure 5e and f).

To validate the sequencing results and understand better the distribution of different pre-rRNA molecules within neurites, we adapted a highly-sensitive non-radioactive Northern Blot method for small RNAs (41) and used it to detect pre-5.8S molecules in neuronal compartments (Figure 6a). We designed a 24 nt long probe conservatively close to the 3’ end of the mature 5.8S that not only would recognize 4 out of the 5 pre-5.8S molecules identified by our pre-rRNAseq, but also longer pre-rRNA species from earlier biogenesis steps. Additionally, we used a probe against the small nucleolar RNA C/D box 104 (SNORD104) as a nuclear control and a probe against the mature 5.8S rRNA as a loading control. We then used compartmentalized chambers to purify RNAs from either the somata+neurite or neurite-only compartment and processed them for Northern Blot. The pre-rRNA probe yielded a double band at around ∼180nt (corresponding to the 179 and 187nt long species identified in our sequencing data), a second band just above 300nt (corresponding to the 272 and 278nt long species identified by sequencing), a third band above the 1000nt marker and some extra signal in the well bottom, most likely representing even larger RNA molecules which did not enter the gel (Figure 6b). Based on our own data and previous studies in humans (30, 42, 43), we interpreted the 180nt and 300nt bands as cytosolic pre-5.8S and the >1000nt band as one of the last nuclear pre-5.8S species. We observed a very strong de-enrichment of the nuclear 1000nt long pre-5.8S in the neurite fraction, which was also observed for SNORD104, as expected (Figure 6b and c) – confirming the purity of our preparation. Both cytosolic pre-5.8S species were, in contrast, highly abundant in the neurite compartment, and even slightly enriched compared to the somata+neurite compartment (Figure 6b and c). Altogether, our data dovetail with the detection of cytosolic RBFs in dendrites and show that cytosolic (but not nuclear) pre-rRNAs are localized to neuronal processes.

**Figure 6.**
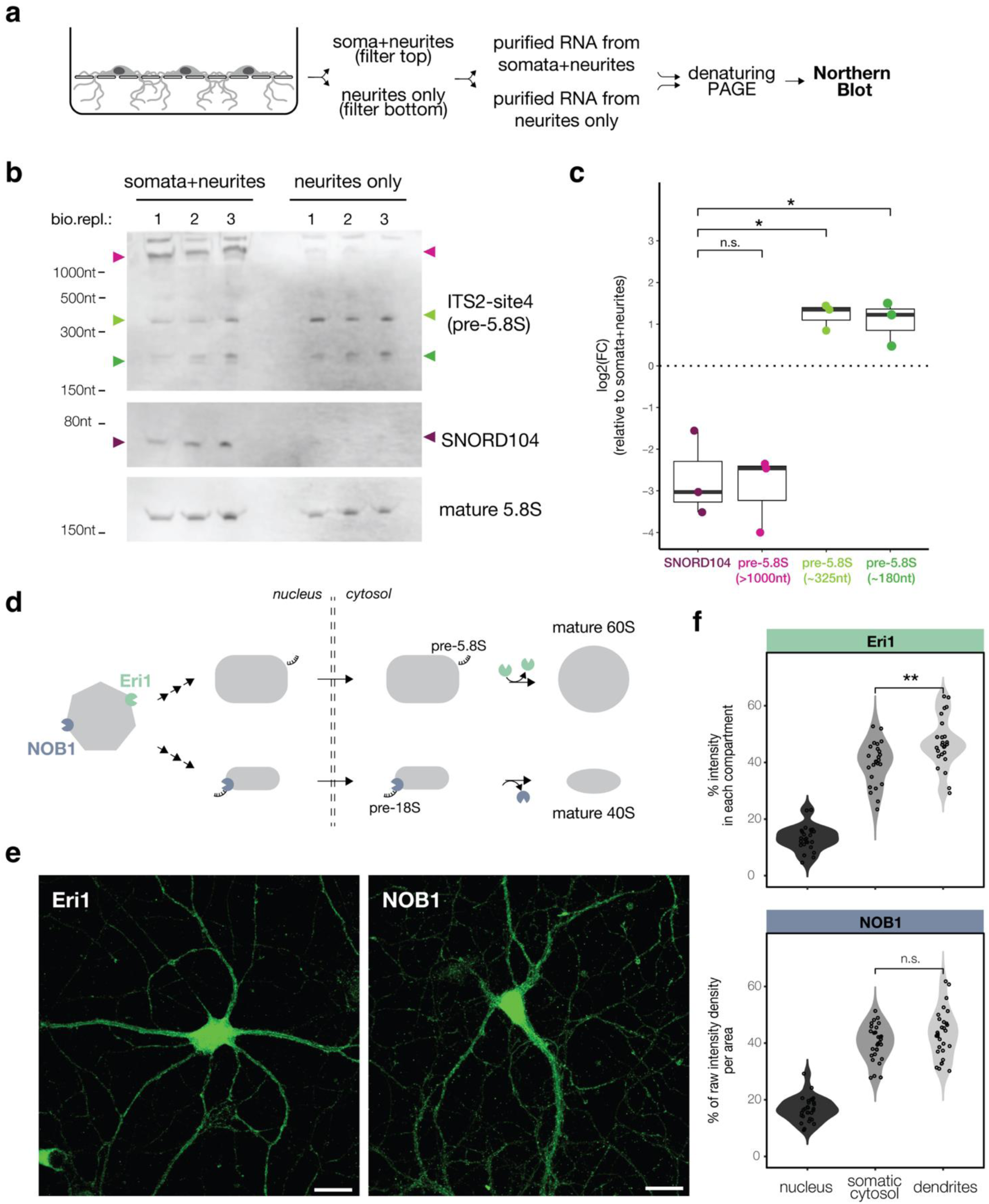
Cytosolic pre-rRNA species and their nucleases are abundant in processes. (**a**) Schematic of the experimental design. RNAs were purified from either subcellular compartment (somata+neurites vs neurites only) and run on a denaturing TBE-Urea gel to separate RNAs of different sizes. RNA molecules were measured by Northern Blot. (**b**) Detection of RNA molecules of interest across subcellular compartments by Northern Blot (three biological replicates). (**c**) Boxplot of the normalized intensity levels of the indicated RNA molecules in the neurite-only compartment (data shown in b). Each dot represents an independent biological replicate. Anova, p = 0.00014; *t*-test between SNORD104 and pre-5.8S (>1000nt), p = 0.78; *t*-test between SNORD104 and pre-5.8S (∼325nt), p = 0.015; *t*-test between SNORD104 and pre-5.8S (∼180nt), p = 0.011. (**d**) Schematic representation of the incorporation and dissociation of the endonuclease NOB1 and the exonuclease Eri1 (respectively in blue and green) during the maturation of the small and large ribosomal subunits (in gray). (**e**) Representative images showing the localization (within individual neurons) of the endogenous nucleases NOB1 and Eri1 (scale bar 25μm). (**f**) Violin plot of the percentage of raw intensity density per area. Each dot represents an individual neuron, from four independent experiments. Pairwise comparisons were performed between compartments of the same individual neuron. For Eri1, Kruskal−Wallis, p = 2.9e−12; *t*-test between somatic cytosol and dendrites, p = 0.0033. For NOB1, Kruskal−Wallis, p = 2.9e−13; *t*-test between somatic cytosol and dendrites, p = 0.2.

### The enzymes for late-stages of rRNA processing are present in neuronal processes

Lastly, we asked whether the machinery necessary to carry-out the late states of rRNA processing is present in neuronal processes and whether distally localized pre-rRNA molecules could undergo local maturation. The endoribonuclease NOB1 is known to stably associate with the Internal Transcribed Spacer ITS1 in the nucleolus and to catalyze the removal of final part of ITS1 (site D/3) in the cytosol, generating the mature 18S rRNA (44, 45) (Figure 6d). The exonuclease Eri1 transiently interacts with the 5’ External Transcribed Spacer ETS in the nucleolus and catalyzes the final 3’ trimming step of the Internal Transcribed Spacer ITS2 to generate the mature 5.8S rRNA in the cytosol (46) (Figure 6d).

We performed immunostaining in cultured hippocampal neurons with antibodies against NOB1 and Eri1. As expected, the antibodies to both nucleases gave a strong signal in the nucleolus. The cytoplasmic signal, however, was not confined to the perinuclear region but distributed throughout the dendrites, present at either similar (NOB1) or slightly higher (Eri1) levels than observed in the perinuclear region (Figure 6e and f). Altogether, these data show that cytosolic pre-rRNAs are localized to dendrites where the machinery is present for their local processing.

## Discussion

Using an array of methods, including both targeted and unbiased strategies, we describe here for the first time the localization of immature ribosomes in the dendrites and/or axons of neurons. Our findings reveal that all constituent elements of cytosolic, but not nuclear, pre-mature ribosomes, including both RNA and protein components, are present in neuronal processes. Contrary to prevailing models which assume that ribosome biogenesis concludes in the perinuclear region, with only fully assembled ribosomes being subsequently transported to distal subcellular compartments, our observations indicate a significant presence of pre-mature ribosomal particles within neuronal processes.

We found that the abundance of immature ribosomes in neuronal processes was substantial and comparable to their levels within the cell body. This observation suggests that they are actively localized, and further studies are clearly needed to dissect transport mechanisms. RNA granules have already been shown to contain translationally incompetent ribosomes, defined by the presence of RPs and rRNAs, but the absence of tRNAs, translation factors like eIF4E and 4G and the inability to incorporate amino acids (17, 47). All of these features would apply also to pre-mature ribosomes. However, additional studies are required to query the presence of RBFs or pre-rRNA species in such granules, and investigate their potential trafficking via microtubules.

Additionally, neurons may localize immature ribosomes in their distal processes also via the Endoplasmic Reticulum (ER). In fact, a recent publication reported a specific subpopulation of 60S ribosomes tethered to the ER of mammalian cell lines and bound by the cytosolic RBFs Lsg1, Ndm3, Zfp622 and eIF6 (48). Interestingly, we also observe these factors at high levels in neuronal processes (Figure 2c). However, their potential association with ER needs to be tested.

We observed that markers of pre-ribosomal particles from the latest steps of ribosome biogenesis tend to be particularly enriched in neuronal processes (axons and dendrites) compared to those associated with earlier steps (Figure 3c-f). We also observed that three RBFs (RSL24D1, Bystin, and RIOK2) were present within synaptic fractions purified from rat cortex. One of these factors, RIOK2, was present in synapses at levels comparable to the integral synaptic scaffolding molecule, PSD-95 (Figure 3e-f). These data suggest that neurons could be poised to complete ribosome maturation very rapidly, for example in response to external or synaptic stimuli. Together with other mechanisms neurons use to reorganize their translational machinery and alter protein synthesis during plasticity (3, 7, 15, 49), our data suggest the local maturation of ribosomes could efficiently boost the local capacity for translation. This potential regulation could be especially relevant in small compartments, such as the dendritic spine, where protein concentrations are tightly regulated and minor changes can have substantial effects. In all our data sets, we consistently observed that the proportion of mature rRNA and RPs is enriched in the cell body, while pre-rRNAs and RBFs tend to be equally distributed throughout the cytosol (including dendrites). Effectively, this results in a higher ratio of immature-to-mature ribosomes specifically in neuronal processes. We propose that the on-demand maturation of immature ribosomes could have a large impact near synapses, resulting in the fine-tuned regulation of the number of ribosomes able to supply a synaptic neighborhood with protein synthesis. The spatial location of these nascent ribosomes as well as their interactions partners could lead to biases in the translation of specific mRNAs.

Finally, we note that the mRNAs encoding RPs that are incorporated during the cytosolic steps of ribosome biogenesis tend to be more abundant in dendrites compared to those of early nuclear steps (Suppl. Figure 3b). We hypothesize that neurons could use the context-dependent translation of RP mRNAs as a mechanism for the rapid local activation of immature ribosomes. Additionally, via the differential incorporation of locally synthesized RPs (25), immature ribosomes might acquire different properties during the last steps of biogenesis that could differentiate them from the local pool of pre-existing ribosomes. Altogether, our data suggests a novel potential mechanism for protein synthesis regulation near synapses.

## Materials and Methods

### Animals

All animals were housed on a 12/12-h light/dark cycle with food and water *ad libitum* until euthanasia.

### Preparation of primary cultured neurons

Dissociated rat hippocampal or cortical neuron cultures were prepared and maintained as described previously (50). Briefly, we dissected hippocampi or cortices from postnatal day 0 to 2 rat pups of either sex (Sprague–Dawley strain; Charles River Laboratories) and dissociated the samples with papain (Sigma). For imaging experiments, hippocampal neurons were plated at a density of 30 × 10^3^ cells/cm^2^ (for antibody staining against endogenous proteins) or 40 × 10^3^ cells/cm^2^ (for transfection experiments) on poly-D-lysine–coated glass-bottom petri dishes (MatTek P35G-1.5-14-C). For biochemical experiments, cortical neurons were plated at a density of 4 × 10^6^ cells/cm^2^ on poly-D-lysine-coated 10 cm dishes or at a density of 9 × 10^6^ cells/cm^2^ on poly-D-lysine-coated 75 mm inserts (compartmentalized chambers; 3.0μm pore size, NEST 726001). One day after plating on the inserts, cells were incubated with 5μM AraC (Sigma C1768) for 2 days, then the AraC was removed by changing the media. Neurons were maintained in a humidified atmosphere at 37 °C and 5% CO2 in growth medium [Neurobasal-A supplemented with B27 and GlutaMAX-I (Life Technologies)] for 14–16 DIV for biochemical experiments or for 25–27 DIV for imaging experiments.

### Immunostaining of primary cultured neurons

Cells were fixed for 20 min in 4% PFA in 4% sucrose in PBS, permeabilized for 15 min in 0.5% Triton-X-100 + blocking buffer and blocked for at least 30 min in blocking buffer (PBS + 4% goat serum). Neurons were incubated for 2 h with primary antibodies (Suppl. Table 2) and, after three washes, for 1 h with secondary antibodies (Suppl. Table 2), all in blocking buffer. After two washes in PBS, cells were stained with DAPI (1:1000 for 2 min) and kept in PBS at 4 °C until imaging. For validation of the compartmentalized chambers (Figure 1a), pieces of the filter were processed as described above, and mounted on a glass slide (ThermoFisher 10417002) with Aqua Poly/mount (Polysciences, 18606). Samples were imaged using a LSM780 confocal microscopy (Zeiss) using a Plan-Apochromat 40x/1.4 Oil DIC M27 or Plan-Apochromat 20x/0.8 M27 objectives. A z-stack was set to cover the entire volume of the neuron, with optical slice thickness set to optimal and number of slices kept equal across conditions of the same experiment. Laser power and detector gain were adjusted to avoid saturated pixels. Imaging conditions were held constant within experiments. Average intensity projections of z-stacks were used for image analysis. For visualization purposes (but not analyses) brightness and contrast were adjusted.

### Image analysis

All image analyses were performed in ImageJ/FIJI (Version 2.1.0/1.53c) with a semi-automated script built in-house. Briefly, the soma and dendritic arbor of individual neurons were manually traced based on the MAP2-positive signal. The nucleus was automatically determined using the DAPI signal. The somatic cytosol area was determined by subtracting the nuclear (DAPI-positive) area from the manually traced somatic area. The raw intensity of the channel of interest (the Dendra2-fluorescent reporter for Figure 3b-d or the antibody staining for Figure 6e-f) was then measured in each area. Additionally, for Figure 2d and e, the background intensity in empty regions was subtracted from the intensity in the neuronal subcellular compartment from the same field of view. Statistical testing and plotting were then performed in RStudio (Version 2023.06.1+524).

### Sample collection from compartmentalized chambers for protein analysis

Both sides of the inserts were washed three times with ice-cold RNase free DPBS (ThermoFisher, 14040-091) supplemented with 100 µg/mL of CHX (Sigma, C7698). The upper compartment (containing somata and neurites) was scraped in 400µL of RNase-free PBS. The lower compartment (containing neurites only) was scraped twice in 100µL of ribosome lysis buffer (20 mM Tris pH 7.4, 150 mM NaCl, 5 mM MgCl2, 24 U/mL TurboDNase, 100 µg/mL cycloheximide, 1% TritonX-100, 1 mM DTT, RNasin(R) Plus RNase inhibitor 200U/mL and 1x cOmplete EDTA-free protease inhibitor). To guarantee sufficient yield from the neurite compartment, each biological replicate consisted of the content of two inserts pooled together. Each sample was then split (10% used for total lysate, 90% for sucrose cushioning). The samples from the upper compartment were centrifuged at 500 × g at 4°C for 5 min, supernatant was discarded and cell pellets were used for downstream processing.

### Sample collection from compartmentalized chambers for RNA analysis

Both sides of the inserts were washed three times with ice-cold RNase free PBS. The upper compartment (containing cell body and neurites) was scraped in 400µL of RNase-free PBS. The lower compartment (containing neurites only) was scraped twice in 100µL of RNase-free PBS. All samples were centrifuged for 5 min at 500 × g at 4 °C. Most of supernatant was discarded, leaving only 200µL of volume. 600µL of Trizol LS solution (ThermoFisher, 10296010) was added to each sample, and mixed well by vortexing. Samples were either immediately used for RNA purification or stored at -80 °C. Samples containing soma+neurites were purified using Direct-zol RNA Miniprep Plus (Zymo Research, R2072) and eluted in 50 µL of RNase free water. Samples containing neurites only were purified using Direct-zol RNA Microprep (Zymo Research, R2062) and eluted in 10µL of RNase free water. In both cases, the manufacturer instructions were followed (including the DNAse I step). The RNA content from two inserts were pooled to have enough material for downstream experiments.

### Total cell lysates

Cells were lysed in 200 µL (upper compartment) or 50 µL (lower compartment) of 8 M urea, 200 mM Tris/HCl [pH 8.4], 4% CHAPS, 1 M NaCl, cOmplete EDTA-free protease inhibitor (Roche, 11873580001), using a pestle. Lysates were sonicated at 4°C for 4 rounds of 30 sec each, and incubated for 10 min with 1 µL of Benzonase (Sigma E1014). After centrifugation for 5 min at 10,000 × g, the supernatant was saved and protein concentration was measured with BCA assay (ThermoFisher, 23252). Samples (between 10 to 60 µg) were then submitted to Mass Spectrometry.

### Sucrose cushion

Cell pellets from the upper compartments were resuspended in 400 µL of ribosome lysis buffer. All samples were pipetted up and down with a 0.4x20mm syringe needle (HSW FINE-JECT) on ice until homogenization was clear. Samples were then centrifuged at 10,000 × g for 10 min at 4°C. Supernatants were loaded on 1 mL sucrose solution (34% sucrose, 20 mM Tris pH 7.4, 150 mM NaCl, 5 mM MgCl2, 1 mM DTT, 100 µg/mL cycloheximide) in a thickwall polycarbonate tube (Beckman, 349622) and centrifuged for 3 hours and 30 min at 4°C at 55000 rpm (367600 × g) with a SW55Ti rotor (acceleration 0, deceleration 7). Ribosome pellet was resuspended in 30 µL of 10 mM HEPES, 120 mM NaCl, 3 mM KCl, 10 mM D-Glucose, 2mM MgSO4 and 2 mM CaCl2, and submitted to Mass Spectrometry

### Western Blot validation of sucrose cushioning

Cells from two 10 cm dishes were washed three times with 10 mL of ice-cold RNase free DPBS (ThermoFisher, 14040-091) and then scraped in 500µL of ice-cold RNase free DPBS supplemented with 100μg/mL cycloheximide. Cells were then pelleted at 500 x *g* for 5 min at 4C and the supernatant discarded. Cell pellets were then resuspended in 500μL ribosome lysis buffer. All samples were pipetted up and down until homogenization was clear with a 0.4x20 mm syringe needle (HSW FINE-JECT) on ice. Samples were then centrifuged at 10,000 x *g* for 10 min at 4C. 200 µL of the supernatants (cleared lysates) were loaded on 1 mL sucrose solution in a thickwall polycarbonate tube (Beckman, 349622) and centrifuged for 30 min or for 3 hours and 30 min at 4C at 55000 rpm (367600 x *g*) with a SW55Ti rotor (acceleration 0, deceleration 7). Ribosome pellet was resuspended in 20 μL of 10 mM HEPES, 120 mM NaCl, 3 mM KCl, 10 mM D-Glucose, 2mM MgSO4 and 2 mM CaCl2, and submitted to Western Blot. 6 µL of the clear lysates (input, 3% of the input total volume) or 10 µL of the cushion samples (50% of the total volume) were mixed with NuPAGE LDS Sample Buffer (ThermoFisher, NP007) and NuPAGE Sample Reducing Agent (ThermoFisher, NP004) to a final concentration of 1x, and then loaded onto a 4% to 12% Bis-Tris NuPAGE gel (ThermoFisher). Gels were transferred using the Trans-Blot Turbo Transfer Pack (Biorad, 1704157) on a polyvinylidene fluoride membrane (Immobilon-FL, IPFL00010 0.45μm pore size). Membranes were stained using Revert 700 Total Protein Stain (LI-COR, 926-11015) to access loading, and then destained according to the manufacturer’s instructions. Immunoblotting was performed with primary and secondary antibodies (see Suppl. Table 2), diluted in Intercept Blocking Buffer (LI-COR 927-70001). 0.1% Tween in PBS was used for washes. Images were acquired using LI-COR Image Studio Lite (version 3.0.30, RRID:SCR_013715). Band intensity was quantified with AzureSpot Pro (v1.0 – 366, Azure Biosystems). Image analysis was then performed in RStudio (Version 2023.06.1+524).

### Synaptosome preparation

For each replicate, one Cortex hemisphere dissected from 8-9 weeks old male Sprague Dawley rats (CD, Charles River Laboratories) after decapitation under deep isoflurane anesthesia was homogenized in a Glass-Douncer in 5 mL SynPer reagent (ThermoFisher) supplemented with Protease Inhibitor Cocktail III (Calbiochem) by 10 careful strokes with a loose and 10 strokes with a tight pestle on ice. An aliquot of the homogenate was set aside and 4 ml of the homogenate were centrifuged 3 times for 10 min at 4°C at 1200 x g to ensure efficient removal of cell debris and nuclei. Subsequently the Synaptoneurosomes (SN) were collected as pellet from 3 mL of the supernatant by centrifugation for 20 min at 15000 x*g* at 4°C and resuspended in 500 µL SynPer reagent with protease inhibitor. Lysates were prepared from homogenate and SN fractions by addition of water, 4xLDS sample buffer (ThermoFisher) and 10x reducing agent (ThermoFisher) to a final concentration of 1ug/ul total protein and 20 ug were separated on NuPAGE 4-12% BIS TRIS 1.5 mm gels (ThermoFisher) with MOPS running buffer and blotted with the Trans-Blot Turbo Transfer system (BioRAD) onto Nitrocellulose membranes. Total protein was visualized by staining with Ponceau S solution (ThermoFisher). Primary antibodies were applied over night at 4°C, IRDye680-coupled secondary antibodies against the respective species were used 1:5000 and applied for 1 hr at room temperature prior to scanning on an Azure Sapphire imager.

### Sample preparation for MS analysis

Proteins were digested using S-Traps according to an adapted version of the suspension trapping protocol described by the manufacturer (ProtiFi, Huntington, NY). Peptides were desalted using C18 StageTips (Rappsilber, Juri, 2007), dried by vacuum centrifugation and stored at -20°C until LC-MS analysis.

### LC-MS/MS Analysis

Dried peptides were reconstituted in 20 µl of 95% H_2_O, 5% acetonitrile (ACN) with 0.1% FA. Peptides were loaded onto a C18-PepMap 100 trapping column (particle size 3 µm, L = 20 mm, ThermoFisher Scientific) and separated on a C18 analytical column with an integrated emitter (particle size = 1.7 µm, ID = 75 µm, L = 50 cm, CoAnn Technologies) using a nano-HPLC (Dionex U3000 RSLCnano). Temperature of the column oven was maintained at 55 °C. Trapping was carried out for 6 min with a flow rate of 6 µL/min using a loading buffer (100% H_2_O, 2% ACN with 0.05% TFA). Peptides were separated by a gradient of water (buffer A: 100% H_2_O and 0.1% FA) and acetonitrile (buffer B: 80% ACN, 20% H_2_O and 0.1% FA) with a constant flow rate of 250nL/min. Peptides were eluted by a non-linear gradient with 120 min active gradient time, as selected for the respective MS method by (51); total run including loading, washing and equilibration was 155 min. Analysis was carried out on a Fusion Lumos mass spectrometer (ThermoFisher Scientific) operated in positive polarity and data independent acquisition (DIA) mode. The DIA method defined MS1 scans followed by 40 DIA scans with optimized segment widths, as published by (51). In brief, the DIA-method had the following settings. Full scan: orbitrap resolution = 120k, AGC target = 125%, mass range = 350-1650 m/z and maximum injection time = 100 ms. DIA scan: activation type: HCD, HCD collision energy = 27%, orbitrap resolution = 30k, AGC target = 2000%, maximum injection time = dynamic. The mass spectrometry proteomics data have been deposited to the ProteomeXchange Consortium via the PRIDE (52) partner repository with the dataset identifier PXD058169.

### MS-data processing

DIA raw files were processed with the open-source software DIA-NN (version 1.8.2 beta 27) using a library-free approach. The predicted library was generated using the in silico FASTA digest (Trypsin/P) option with the UniProtKB database (Proteome_ID: UP000002494) for *Rattus norvegicus*. Deep learning-based spectra- and RT-prediction was enabled. The covered peptide length range was set to 7-30 amino acids, missed cleavages to 2 and precursor charge range to 1-5. Methionine oxidation and protein N-terminal acetylation were set as variable modifications. Cysteine carbamidomethylation was selected as a fixed modification. The maximum number of variable modifications per peptide was limited to 1. According to most of DIA-NN’s default settings, MS1 and MS2 accuracies as well as scan-windows were set to 0, isotopologues were enabled, while match-between-runs and shared spectra were disabled. Protein inference was performed using genes with the heuristic protein inference option enabled. The neural network classifier was set to single-pass mode and the quantification strategy was selected as “QuantUMS (high precision)”. The cross-run normalization was set to “RT-dependent”, the library generation to “smart profiling”, the speed and Ram usage to “optimal results”.

### MS Post-processing and statistical analysis

The DIA-NN report table was imported in the statistical computing software R. The entries were filtered for 1% FDR at protein (Global.PG.Q.Value and PG.Q.Value) and precursor level (Q.Value) as well as for unique peptides (Proteotypic) before performing the protein roll-up of the precursor intensities using the diann_maxlfq function provided by the DIA-NN R package (https://github.com/vdemichev/DiaNN). For further analysis, protein intensities were log2-scaled and normalized using median-centering. For the differential expression analysis, only genes with at least 7 observations across replicates were considered.

GO overrepresentation analysis was performed using the ClusterProfiler R package (https://github.com/YuLab-SMU/clusterProfiler). In brief, gene lists of regulated proteins were compared to a custom background dataset containing all genes identified in the experiment. The analysis was then carried out using gene symbols, the ontology setting “cellular compartment”, Benjamini-Hochberg FDR correction and an FDR cut-off of 1%. Redundant GO terms were simplified according to the adjusted p-value with a redundancy cut-off of 0.7 before merging the enrichment results for visualization of the top-25 terms per condition.

### Molecular Cloning

Constructs containing Dendra2 fused to a Ribosome Biogenesis Factor (Pescadillo, DIMT1 or RIOK2) or Ribosomal Protein RPS14 were cloned using Gibson Assembly (NEB, E5510) and standard cloning techniques. In each case, the insert for Dendra2 was obtained as a gBlocks Gene Fragment (IDT), and the CAG vector backbone was derived from CRY2-GFP-homer1c (Addgene plasmid #89442, a gift from Matthew Kennedy) by XhoI and EcoRV digestion to excise CRY2-GFP-homer1c. The unique inserts for the DIMT1 and RIOK2 constructs were amplified by PCR from DIMT1_pLX307 (Addgene plasmid #98330, a gift from William Hahn & Sefi Rosenbluh), and pDONR223-RIOK2 (Addgene plasmid #23535, a gift from William Hahn & David Root), respectively. The unique insert for RPS14 (sequence provided below) was amplified by PCR from a pcDNA3.1 vector (Invitrogen, V79020) containing the RPS14 open reading frame. The unique insert for Pescadillo was obtained as a gBlocks Gene Fragment (IDT).

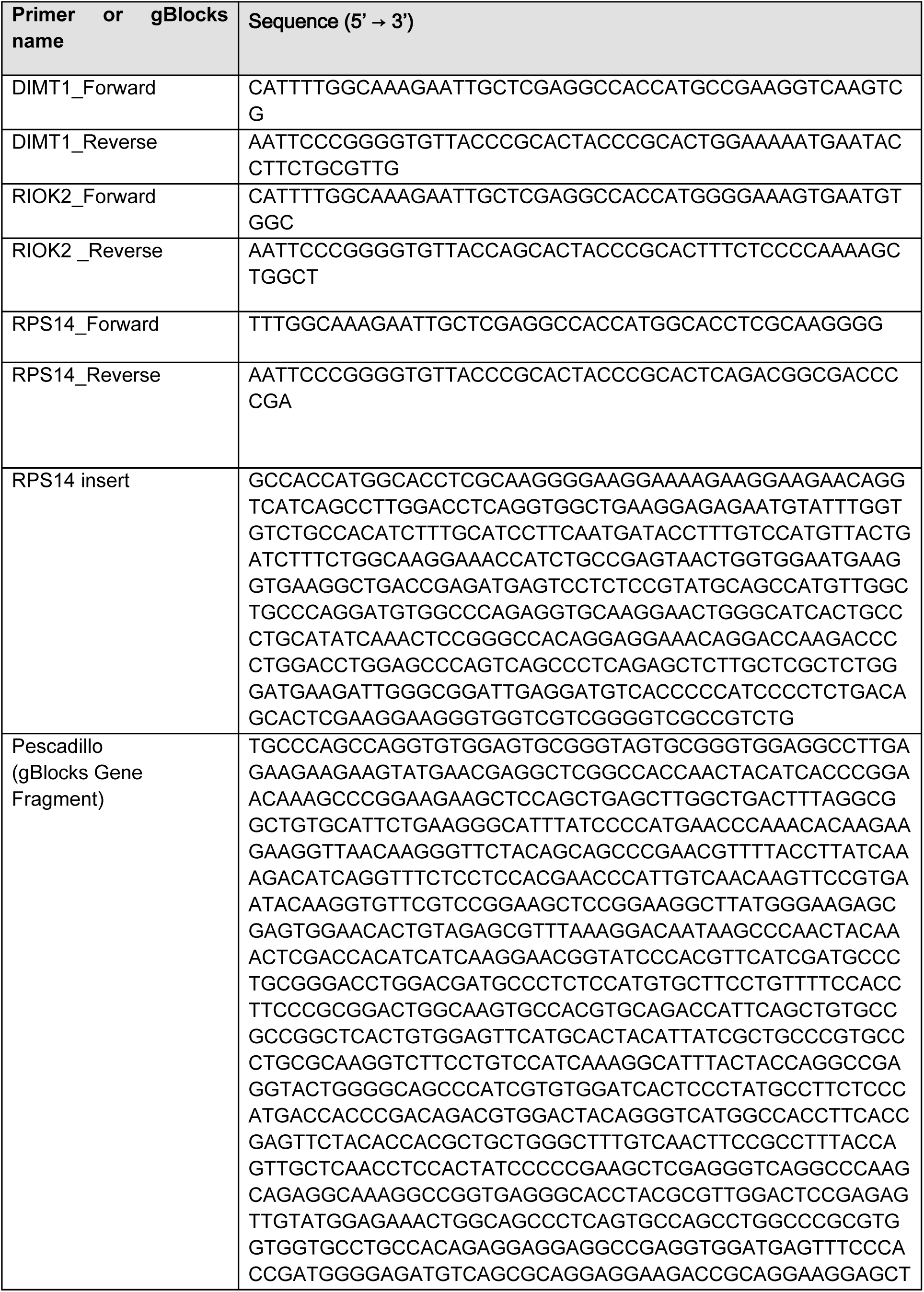
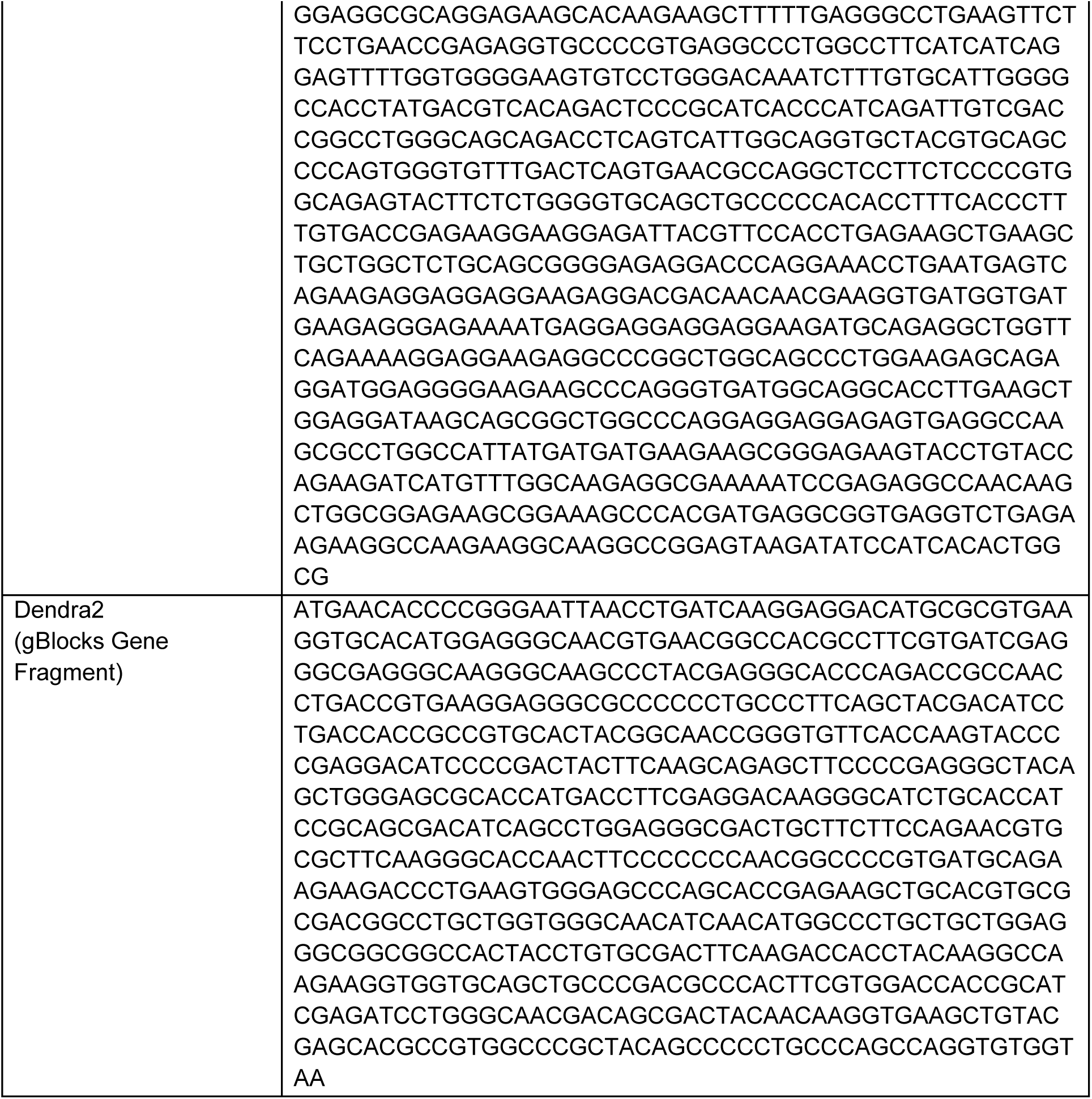

### Gel electrophoresis for Northern Blot

Samples (1µg of RNA) were added to a volume of the same size of 2X RNA Loading Dye (NEB, B0363A). RNAs were denatured at 70 °C for 10 min and returned to ice before being loaded on a denaturing Novex 10% TBE-Urea Gels (ThermoFisher, EC6875BOX) in 1X TBE running buffer at 200V for 90 minutes. Total RNA was visualized with SYBR Gold Nucleic Acid Gel Stain (ThermoFisher, S11494).

### Northern Blot transfer and crosslink

Northern Blot procedure was performed as previously described (41). Briefly, RNA was transferred to an Ambion BrightStar-Plus membrane (ThermoFisher, AM10100) at 30V for 2 hours at 4 °C, using a semi-dry transfer cell (BioRad, 170-3957). Membrane was washed in 2x SSC and then placed on a Whatmann paper soaked in 0.753 grams of 1-ethyl-3-(3-dimethylaminopropyl) carbodiimide (EDC; Sigma, 7750) dissolved in 1.25 M 1-Methylimidazole (Sigma, M50834) pH 8.0. To facilitate RNA–membrane cross-linking, the membrane was incubated at 60 °C for 2 hours. Residual cross-linking reagents were removed by thoroughly rinsing the membrane with distilled water.

### Probes generation for Northern Blot

Probes with the following sequence were purchased as DNA oligos from IDT:

- ITS2-site4 (GGG+CGGCG+ATTGA+TCGTC+AAGCGA, where + indicates LNA base)
- SNORD104 (GATTCGCATCACCCGGATCAGCAGTCTAACGC)
- mature 5.8S (GGCCGCAAGTGCGTTCGAAGTGTCGATGAT).

To generate DIG containing probes, the DIG Oligo tailing kit (Roche, 3353583910) was used. Briefly, 2µL containing 50pmol of probes were mixed with RNase free water (7µL), reaction buffer (4µL), CoCl2-solution (4µL), DIG-dUTP solution (1µL), dATP solution (1µL) 400 U Terminal transferase (1µL), and incubate for 15 min at 37 °C. The reaction was stopped by adding 2µL 0.2M EDTA (pH 8.0).

### Northern Blot hybridization and detection

Membranes were pre-hybridized in 10 mL of ULTRAhyb Ultrasensitive Hybridization Buffer (ThermoFisher, AM8670) for at least 30 min at 42 °C. Probes were heated at 95 °C for 1 min and then added to the prehybridized blot at a final concentration of 0.5nM overnight (for ITS2-site4 and SNORD104) or 0.025nM for 30 min (for mature 5.8S). Blots were washed twice for 5 min at 42 °C in 2X SSC, 0.1% SDS, twice for 15 min at 42 °C in 0.1X SSC, 0.1% SDS, and once for 5 min at room temperature in 1x DIG Washing Buffer from the DIG Wash and Block Buffer Set (Roche, 11585762001). Membranes were incubated for 3 hours at room temperature in 50 mL of 1x DIG Blocking buffer in 1x Maleic Acid Buffer. Anti-Digoxigenin-AP, Fab fragments (Roche, 11 093 274 910) were then added (1:10,000) and the membranes were incubated at room temperature for 30 min. Membranes were then washed twice for 15 min in 1x DIG Washing Buffer, and once for 5 min in 1x DIG detection buffer. CSPD (Roche, 11 655 884 001) was then diluted 1:100 in 1x DIG detection buffer and applied to the membrane. Blots were inserted in a Blot Development folder (Azure Biosystems, AC2126) and incubated protected from light at room temperature for 5 min, and 37 °C for 10 min. Membranes were kept protected from light at room temperature for at least 30 minutes prior to imaging. Stripping was performed following the protocol available online from the Drummond lab. Briefly, membranes were incubated twice for 15 min in 50 mL of boiling 0.1X SSC, 0.5% SDS, before starting with the hybridization and detection steps for the new probe.

### Northern Blot analysis

Image acquisition was performed at the Azure using the chemiluminescence program. Band intensity was quantified with AzureSpot Pro (v1.0 – 366, Azure Biosystems). Image analysis was then performed in R.

### Gel electrophoresis for pre-rRNA sequencing

Purified RNA samples were mixed with an equal volume of 2X Novex TBE-Urea Sample Buffer (ThermoFisher, LC6876). Samples were denatured at 80 °C for 90 sec and returned to ice before being loaded on a denaturing Novex 10% TBE-Urea Gels (ThermoFisher, EC6875BOX) in 1X TBE running buffer at 200V for 50 minutes. Total RNA was visualized with SYBR Gold Nucleic Acid Gel Stain (ThermoFisher, S11494). Gel bands in the range of 160nt to 400nt were excised and crushed with disposable RNase-free pestles (FisherScientific, 13236679). The crushed gels had the addition of 100µL gel elution buffer composed of 0.3M sodium acetate pH = 4.5, 0.25% SDS, 1mM EDTA pH = 8.0, as previously described (53). RNA was eluted for 10 minutes at 65°C and shaking at 1400rpm in a thermal mixer. Gel pieces were removed with Spin-X centrifuge tube filter 0,22µm (Costar, Corning 8169). RNA was purified by using the RNA Clean & Concentrator-5 Kit (Zymo Research, R1013) and eluted in6µL of RNase-free water. The yield of RNA was quantified using Qubit HS-RNA (ThermoFisher, Q32855). Samples were either processed immediately for library prep or stored at -80 °C.

### RNA library prep

For end repair, 30 ng of RNA was mixed with 0.2 U/µL T4 PNK (NEB, M0201), 1.2 U/µL SUPERase inhibitors (ThermoFisher, AM2694) in 2X T4 RNA Ligase Buffer (NEB, B0216L), and incubated at 37 °C. for 30 min and at 65 °C for 20 min, before returning to ice. For 3’ adapter ligation, samples were then incubated at room temperature for 3 hours under constant motion, in 1 µM Adenylated Fusion Oligo (Table 5. 9), 10 U/µL Truncated KQ Rnl2 (NEB, M0373) in 25% PEG 8000. Unligated adapters were digested by adding 2.2 µL of a mixture of equal volume of Deadenylase (NEB, M0331S) and Exonuclease VIII (NEB, M0545S) to the samples at 30 °C for 15 min for deadenylation, followed by 37 °C for 30 min for digestion, and then at 75 °C for 10 min to inactivate the enzymes. For Reverse Transcription (RT) Primer Activation, samples were mixed with 0.25 U/µL RNase HII (NEB, M0288) in CutSmart buffer (NEB, B7204) and incubated at 37 °C for 30 min, and then at 75 °C for 10 min to inactivate the enzymes. Reverse Transcription was performed by incubating samples at 42 °C for 16 hours, with 10 U/µL Induro RT (NEB, M0681), 0.5 U/µL SUPERase inhibitors (ThermoFisher, AM2694), 1 mM dNTPs, 5mM DTT in 1x NEBNext First-strand buffer (NEB, E7421AVIAL). To digest and inactive the reverse transcriptase, samples were incubated with 16 U/µL Proteinase K (NEB, P8107S) solution at 25 °C for 10 min and then at 75 °C for 5 min. To digest RNA, samples were incubated with 0.11 U/µL RNase H (NEB, M0297) and 1.1 U/µL RNase If (NEB, M0243) at 37 °C for 30 min and then at 70 °C for 30 min. Unligated adapters were digested by adding 1 U/µL Exonuclease III (NEB, M0206) in CutSmart buffer (NEB, B7204) to the samples at 30 °C for 15 min for deadenylation, followed by 37 °C for 30 min for digestion, and then at 70 °C for 30 min to inactivate the enzymes. cDNA polyadenylation was performed by adding 2 U/µL Terminal Transferase (NEB, M0315), 3 mM dATP and incubating the samples at 37 °C for 1 min and at 75 °C for 20 min. Uracil DNA digestion was performed by adding 0.02 U/µL Thermolabile USER II (NEB, M5508S), 0.1 U/µL rSAP (NEB, M0371), 1 µM 2nd Strand_OligoT (Table 5. 9), and incubating the samples at 37 °C for 30 min and at 65 °C for 10 min. The second strand was synthesized by incubating samples with 1 U/µL Large Klenow Fragment DNA Pol (NEB, M0210M), 2 mM dNTP mix at 25 °C for 60 min and at 75 °C for 20 min. Samples were stored at 4 °C overnight. cDNA amplification was performed by adding 1 µM smRNA PCR1 universal primer and 1 µM smRNA PCR_iX index primer (Step III) (Table 5. 9) and 1x KAPA HiFi HotStart Ready Mix (Roche, 09420398001) to the ⅓ (30 µL) of 2nd strand synthesis reaction. Initial denaturation occurred at 98 °C for 30 sec, followed by 19 cycles consisting of a denaturation step at 98 °C for 5 sec, an annealing step at 60 °C for 10 sec, and an extension step at 72 °C for 30 sec. This was followed by a final extension at 72 °C for 5 min.

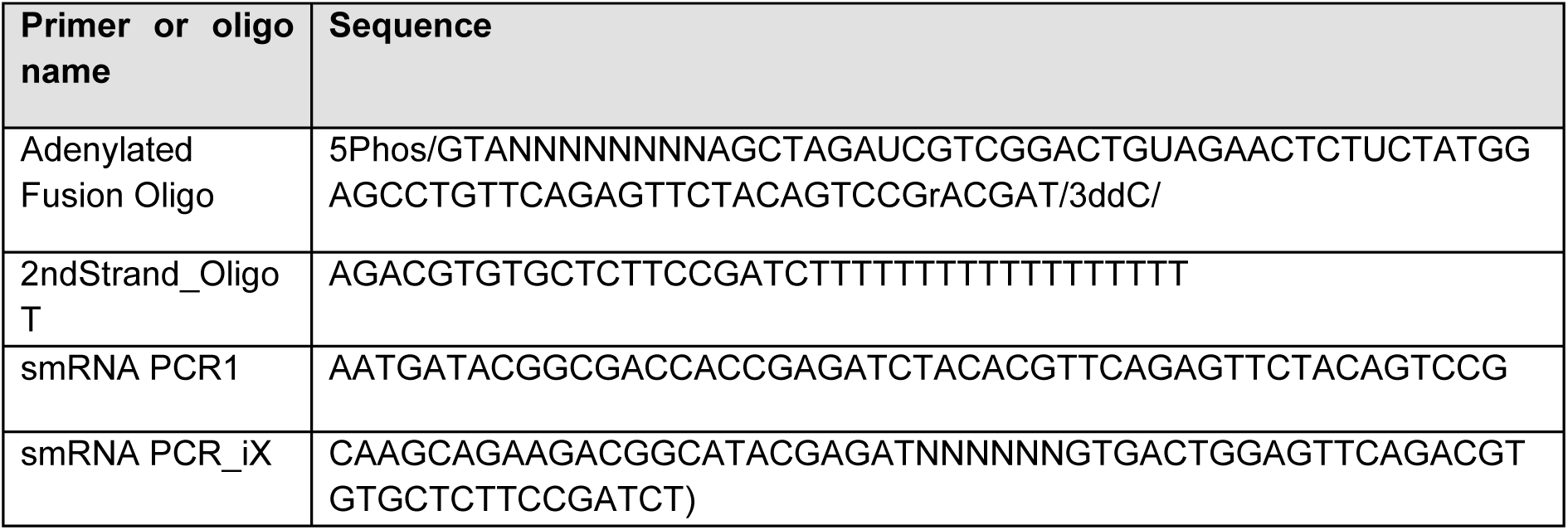

PCR products were purified with Magnetic DNA purification beads (AMPure XP, Beckman Coulter, A63881) according to the manufacturer’s instructions and eluted in 23 µL Buffer EB (Qiagen, 19086). DNA samples were mixed with 6X Gel loading dye (NEB, B7024S) and ran on a 20% TBE gel at 100V for 5 hours. Gels were visualized with SYBR Gold Nucleic Acid Gel Stain (ThermoFisher, S11494). Gel bands in the range of 200 nt to 300 nt were excised and crushed with disposable RNase free pestles. T 20µL of water was added to the crushed gels before being incubated at 4°C for at least 10 hours. Gel pieces were removed with Spin-X centrifuge tube filters (Costar, 8169) and DNA was measured using Qubit dsDNA HS kit (ThermoFisher, Q32854) and the HS DNA Bioanalyzer Assay (Agilent, 5067).

Sequencing was performed using the Illumina NextSeq 2000 system, operating in single-end read mode. This involved a P5 read1 cDNA sequence over 100 cycles and an I7 sample index over 6 cycles, both in the forward direction. Read1 commenced with a 5bp index, an 8bp UMI followed, and a 3 bases anchor (ATG).

### Analysis

For data demultiplexing, we utilized bcl2fastq with a unique base mask (--use-base-mask N5Y*,I6) to exclude the 5bp integrated index from the cDNA’s 5’end. The reads underwent cleaning to remove adapters and low-quality segments using [fastp](https://github.com/OpenGene/fastp). In this process, the UMI was trimmed and added to the header, and the 3 anchor bases were disregarded. The resulting clean reads began with either 3 or 2 base charging states.

Our analysis included preparing a Rat transcriptome reference using the latest rn7 annotation, incorporating the labeling format: <transcript_id>|<gene_symbol>|<rna_type>. Reads were aligned using [bowtie2](https://github.com/BenLangmead/bowtie2), reducing the local alignment’s anchor size to 15. This adjustment was based on the premise that modifications should be detectable within a 15-base window. The alignment parameters were set as follows: ‘-D 20 -R 3 -N 0 -L 15 -i S,1,0.50’.

For gene count tabulation, we developed a custom script. The DESeq2 package facilitated quality control through PCA plots and differential expression testing., All pipeline code is accessible in the [trnatools] (https://gitlab.mpcdf.mpg.de/mpibr/schu/trnatools) repository.

**Table 1.** Manually curated list of Ribosome Biogenesis Factors. List of rat ribosome biogenesis factors, compiled from the literature. The following publications were particularly instrumental for the curation of this list: Bohnsack and Bohnsack, 2019 (54); Dörner et al., 2023 (32); Nerurkar et al., 2020 (38); Prattes et al., 2019 (55). We indicate here the number of biological replicates of total lysates from either compartment (total replicates = 5) where the corresponding gene was identified. Protein intensity values of each replicate are shown in Suppl. Table 2. Additionally, we note that some of these RBFs were previously identified in axonal preparations from mice (Poulopoulos et al., 2019 (23)), rats (Chuang et al., 2018 (56)) and frogs (Cagnetta et al., 2018 (57), Shigeoka et al., 2019 (26)). Abbreviations: small ribosomal subunit (40S), large ribosomal subunit (60S), Nucleus (N), Nucleolus (Nu), Nucleoplasm (NP), Cytoplasm (C).

| Gene name | Ribosomal subunit involved in | Location of first interaction | Location of last interaction | Strength of interaction | Known extra-ribosome biogenesis function? | Number of Soma replicates with gene detected | Number of Neurites replicates with gene detected | Identified in other publications |
| --- | --- | --- | --- | --- | --- | --- | --- | --- |
| Abce1 | 40S | C | C | transient | yes | 5 | 5 | Poulopoulos et al., 2019; Cagnetta et al., 2018; Shigeoka et al., 2019 |
| Bop1 | 60S | Nu | NP | stable | no | 5 | 3 |  |
| Bysl | 40S | N | C | stable | no | 5 | 5 |  |
| Csnk1d | 40S | N | C | stable | yes | 5 | 5 |  |
| Csnk1e | 40S | N | C | stable | yes | 5 | 5 | Chuang et al., 2018 |
| Ddx18 | 60S | Nu | Nu | stable | yes | 5 | 5 |  |
| Dimt1 | 40S | N | C | stable | no | 5 | 0 |  |
| Dusp12 | 60S | N | C | stable | yes | 5 | 5 |  |
| Eif6 | 60S | N | C | stable | yes | 5 | 5 | Chuang et al., 2018; Cagnetta et al., 2018 |
| Fbl | 40S | Nu | Nu | stable | yes | 5 | 5 | Poulopoulos et al., 2019; Chuang et al., 2018 |
| Ftsj3 | 40S | Nu | Nu | stable | yes | 5 | 0 | Chuang et al., 2018 |
| Gtpbp4 | 60S | NP | C | stable | no | 5 | 0 | Shigeoka et al., 2019 |
| Gtpbp4 | 60S | NP | C | stable | no | 5 | 5 | Shigeoka et al., 2019 |
| Llph | 60S | Nu | C | stable | yes | 5 | 0 |  |
| Lsg1 | 60S | C | C | transient | no | 5 | 5 |  |
| Ltv1 | 40S | NP | C | stable | no | 5 | 4 |  |
| Mak16 | 60S | Nu | NP | transient | no | 5 | 1 |  |
| Mdn1 | 60S | Nu | NP | stable | no | 5 | 0 |  |
| Mdn1 | 60S | Nu | NP | stable | no | 5 | 1 |  |
| Mdn1 | 60S | Nu | NP | stable | no | 5 | 3 |  |
| Mphosph10 | 40S | Nu | N | stable | no | 5 | 2 |  |
| Mrt4 | 60S | N | C | stable | no | 5 | 5 |  |
| Nifk | 60S | Nu | N | stable | no | 5 | 1 | Chuang et al., 2018 |
| Nle1 | 60S | NP | C | stable | yes | 5 | 5 |  |
| Nmd3 | 60S | N | C | stable | no | 5 | 5 |  |
| Nob1 | 40S | N | C | stable | no | 5 | 5 |  |
| Nop53 | 60S | Nu | Nu | stable | yes | 5 | 0 |  |
| Nsa2 | 60S | Nu | C | stable | no | 5 | 5 |  |
| Nvl | 60S | Nu | NP | transient | yes | 5 | 0 |  |
| Pdcd11 | both | Nu | N | stable | no | 5 | 5 |  |
| Pelo | 40S | C | C | transient | yes | 5 | 5 | Chuang et al., 2018 |
| Pes1 | 60S | Nu | NP | stable | no | 5 | 5 | Chuang et al., 2018 |
| Pno1 | 40S | N | C | stable | no | 5 | 5 |  |
| Pno1 | 40S | N | C | stable | no | 1 | 0 |  |
| Rae1 | 60S | N | C | stable | yes | 5 | 5 |  |
| Rbm28 | 60S | Nu | Nu | stable | yes | 5 | 1 |  |
| Riok1 | 40S | C | C | stable | yes | 5 | 5 |  |
| Riok2 | 40S | N | C | stable | no | 5 | 5 |  |
| Riok3 | 40S | C | C | NA | yes | 5 | 5 |  |
| Rpf1 | 60S | Nu | NP | transient | no | 4 | 0 |  |
| Rpf1 | 60S | Nu | NP | transient | no | 5 | 0 |  |
| Rrp1 | 60S | Nu | NP | transient | no | 5 | 0 |  |
| Rrp12 | 40S | Nu | C | stable | no | 5 | 5 |  |
| Rsl24d1 | 60S | Nu | C | stable | no | 5 | 0 |  |
| Sbds | 60S | C | C | transient | no | 5 | 5 |  |
| Spata5 | 60S | C | C | transient | yes | 5 | 5 |  |
| Spata5l1 | 60S | C | C | transient | yes | 4 | 4 |  |
| Tsr1 | 40S | NP | C | stable | no | 5 | 5 |  |
| Wdr12 | 60S | Nu | NP | stable | no | 5 | 5 |  |
| Wdr37 | 60S | Nu | NP | stable | yes | 5 | 5 |  |
| Wdr74 | 60S | Nu | Nu | stable | yes | 5 | 1 |  |
| Xrn1 | 40S | C | C | transient | yes | 5 | 5 |  |
| Zfp593 | 60S | Nu | C | stable | yes | 5 | 1 |  |
| Zfp622 | 60S | C | C | stable | yes | 5 | 5 |  |

**Table 2.** List of antibodies.

| Target | Provider | Identifier | Dilution |
| --- | --- | --- | --- |
| Bystin | Kutay lab |  | 1:2500 (WB) |
| Eri1 | abcam | ab236945 | 1:100 (IF) |
| GFAP | abcam | ab7260 | 1:2000 (IF) |
| MAP2 | SYSY | 188004 | 1:2000 (IF) |
| Nob1 | ThermoFisher | A304-680A-T | 1:100 (IF) |
| Nucleolin | abcam | ab22758 | 1:2000 (WB) |
| PSD95 | ThermoFisher |  | 1:1000 (WB) |
| RIOK2 | ThermoFisher | H00055781-B01P | 1:200 (WB) |
| RPS11 | Bethyl | A303-936A | 1:2500 (WB) |
| Rsl24d1 | Santa Cruz<br>Biotechnology | sc-100840 | 1:200 (WB) |
| Goat anti-guinea pig<br>Dylight405 | Jackson<br>ImmunoResearch | 106-475-003 | 1:1000 (IF) |
| Goat anti-guinea pig-Alexa488 | ThermoFisher | A11073 | 1:1000 (IF) |
| Goat anti-mouse-Alexa594 | ThermoFisher | A11005 | 1:1000 (IF) |
| Goat anti-mouse-Alexa488 | ThermoFisher | A11001 | 1:1000 (IF) |
| Goat anti-rabbit-Alexa594 | ThermoFisher | A11037 | 1:1000 (IF) |
| Goat anti- rabbit-<br>Alexa488 | ThermoFisher | A11008 | 1:1000 (IF) |
| Goat anti-rabbit IRDye 680 | LICOR | 926-68071 | 1:5000 (WB) |
| Goat anti-mouse IgG IRDye 680 | LICOR | 926-68020 | 1:5000 (WB) |
| Goat anti-mouse IRDye 800 | LICOR | 926-32210 | 1:5000 (WB) |
| Goat anti-rabbit IRDye 800 | LICOR | 926-32211 | 1:5000 (WB) |

## Acknowledgments

We thank I. Bartnik, N. Fuerst, and C. Thum for the preparation of primary cell cultures, Nassim-Assir B. for hippocampal slice preparation, Dr. S. Junek and Dr. C. Bohnstaedt for assistance with image acquisition, Dr. A. Schwarz and Dr. B.A. Lucas for their dedicated efforts in the 2D cryo-electron microscopy analyses. We are grateful to all members of the Schuman lab. We thank Prof. Dr. Ulrike Kutay for providing the antibody against Bystin/Enp1.

## Author Contributions

Conceptualization: CMF, EMS

Investigation: CMF, AS, AMB, GT, KD, EML, EC, STD, NK, JDL

Formal analysis: CMF, KD, GT

Visualization: CMF, KD

Supervision: EMS, AH

Writing – original draft: CMF, EMS

Writing – review & editing: CMF, AS, AMB, GT, KD, EML, EC, STD, NK, AH, JDL, EMS.

## Classification

Biological Sciences, Cell Biology.

