## Supplemental Figures for "Neuronal processes contain the essential components for the late steps of ribosome biogenesis"

**Supplementary Tables:**

- Suppl. Table 1: manually curated list of translation-related genes (used as input to Fig. 1e)
- Suppl. Table 2: Protein Intensity values of Ribosome Biogenesis Factors identified in total lysates across neuronal subcellular compartments (related to Table 1)
- Suppl. Table 3: Protein Intensity values of all detected proteins in the total lysates and sucrose cushion across neuronal subcellular compartments

**Source Data:**

- Source Data for Figure 1: Differential expression analysis between compartments (total lysate).
- Source Data for Figure 4: Differential expression analysis between total lysate and sucrose cushion from soma+neurites compartment.
- Source Data for Suppl. Figure 3b: RP mRNA abundance per 10μm$^2$ of dendrite (as measured in Fusco et al. 2021) according to the order and site of RP incorporation during ribosome biogenesis.

**Supplementary Figures:**

- Suppl. Figure 1. Proteomic characterization of neuronal subcellular compartments.
- Suppl. Figure 2. Proteomic characterization of sucrose cushioning from different neuronal subcellular compartments.
- Suppl. Figure 3. Customized pre-rRNA-seq pipeline and RP mRNAs localization in dendrites.

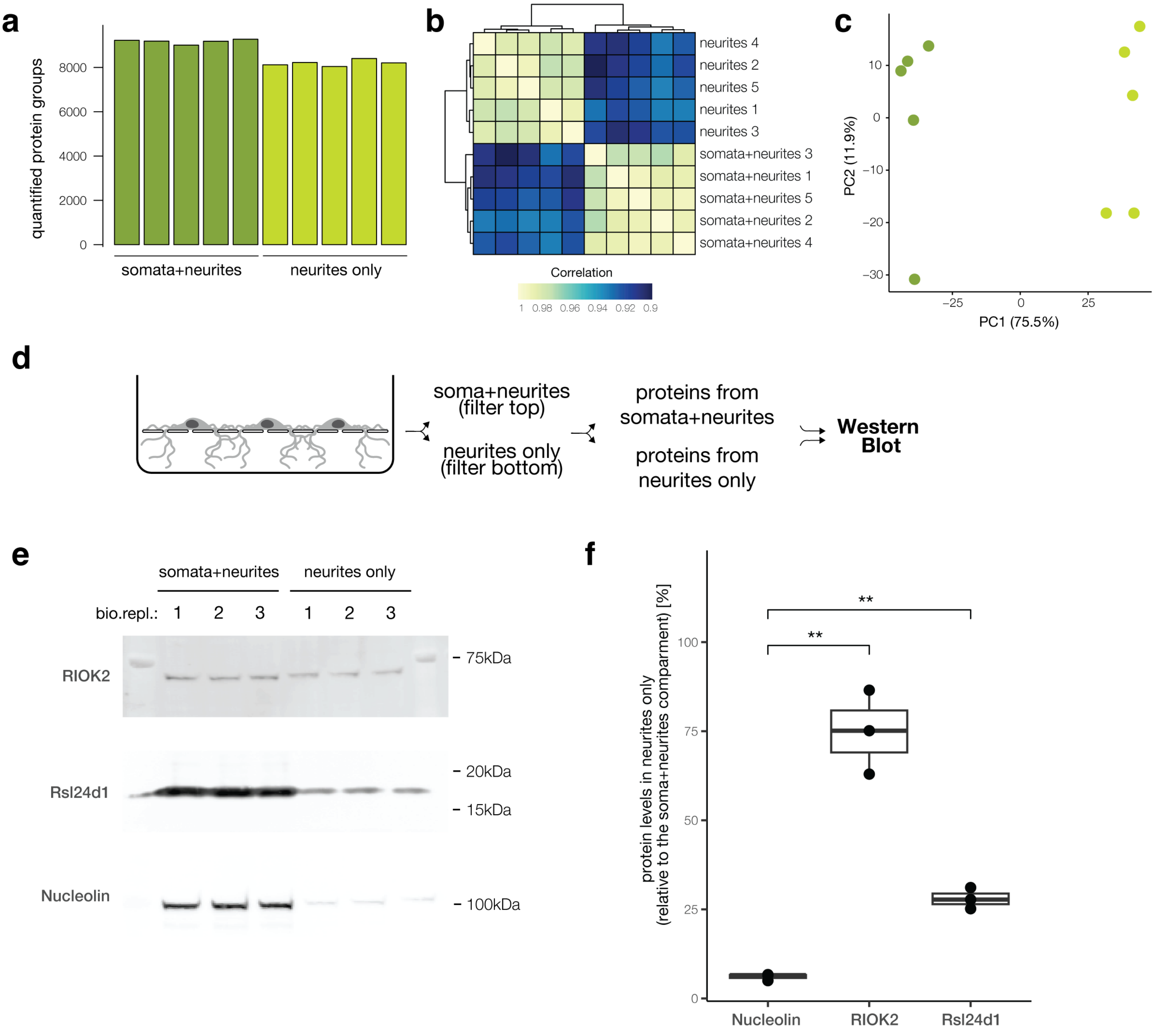
a
quantified protein groups
8000
6000
4000
2000
0
somata+neurites
neurites only
b
neurites 4
neurites 2
neurites 5
neurites 1
neurites 3
somata+neurites 3
somata+neurites 1
somata+neurites 5
somata+neurites 2
somata+neurites 4
Correlation
1 0.98 0.96 0.94 0.92 0.9
c
PC2 (11.9%)
PC1 (75.5%)
10
0
−10
−20
−30
−25
0
25
d
soma+neurites
(filter top)
neurites only
(filter bottom)
proteins from
somata+neurites
proteins from
neurites only
Western
Blot
e
somata+neurites
neurites only
bio.repl.: 1 2 3 1 2 3
RIOK2
Rsl24d1
Nucleolin
75kDa
20kDa
15kDa
100kDa
f
protein levels in neurites only
(relative to the soma+neurites comparment) [%]
100
75
50
25
0
**
**
Nucleolin
RIOK2
Rsl24d1

**Suppl. Fig. 1**

**Suppl. Figure 1. Proteomic characterization of neuronal subcellular compartments.** (**a**) Bar plot of the number of quantified protein groups in the total lysate across biological replicates of each subcellular compartment. The bars related to the somata+neurites compartment are colored in dark green, the ones for neurites only in light green. (**b**) Hierarchical clustering of biological replicates according to protein levels in lysates from somata+neurites or neurites compartments. Cells are color coded according to Pearson correlation coefficients. (**c**) PCA analysis showing similarities across protein levels in lysates from somata+neurites (dark green) or neurites only (light green) compartments. The variability explained by each Principal Component (PC) is shown in brackets. (**d**) Schematic of the experimental design. Proteins were purified from either subcellular compartment (somata+neurites vs neurites only) and measured by Western Blot. (**e**) Detection of protein of interest across subcellular compartments by Western Blot (three biological replicates). (**f**) Boxplot of the intensity levels of the indicated proteins in the neurite-only compartment, normalized to the soma+neurites compartment (data shown in f). Each dot represents an independent biological replicate. Anova, $p = 5.7e{-}05$; *t*-test between Nucleolin and RIOK2, $p = 0.0093$; *t*-test between Nucleolin and Rsl24d1, $p = 0.0033$.

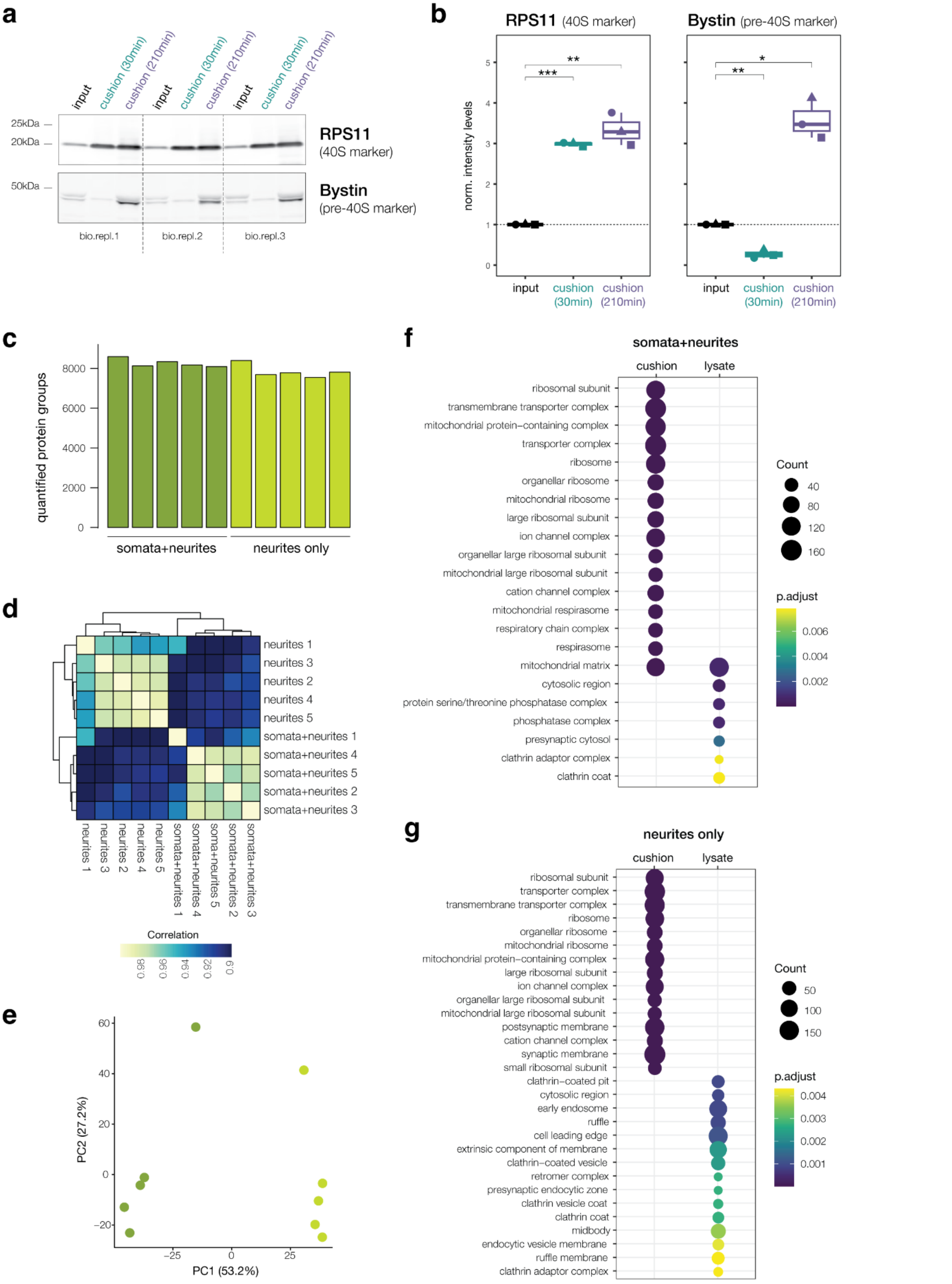
a
input
cushion (30min)
cushion (210min)
25kDa
20kDa
50kDa
RPS11
(40S marker)
Bystin
(pre-40S marker)
bio.repl.1
bio.repl.2
bio.repl.3
b
RPS11 (40S marker)
Bystin (pre-40S marker)
norm. intensity levels
***
**
*
input
cushion (30min)
cushion (210min)
c
quantified protein groups
somata+neurites
neurites only
d
neurites 1
neurites 3
neurites 2
neurites 4
neurites 5
somata+neurites 1
somata+neurites 4
somata+neurites 5
somata+neurites 2
somata+neurites 3
Correlation
e
PC2 (27.2%)
PC1 (53.2%)
f
somata+neurites
cushion
lysate
ribosomal subunit
transmembrane transporter complex
mitochondrial protein−containing complex
transporter complex
ribosome
organellar ribosome
mitochondrial ribosome
large ribosomal subunit
ion channel complex
organellar large ribosomal subunit
mitochondrial large ribosomal subunit
cation channel complex
mitochondrial respirasome
respiratory chain complex
respirasome
mitochondrial matrix
cytosolic region
protein serine/threonine phosphatase complex
phosphatase complex
presynaptic cytosol
clathrin adaptor complex
clathrin coat
Count
p.adjust
g
neurites only
cushion
lysate
ribosomal subunit
transporter complex
transmembrane transporter complex
ribosome
organellar ribosome
mitochondrial ribosome
mitochondrial protein−containing complex
large ribosomal subunit
ion channel complex
organellar large ribosomal subunit
mitochondrial large ribosomal subunit
postsynaptic membrane
cation channel complex
synaptic membrane
small ribosomal subunit
clathrin−coated pit
cytosolic region
early endosome
ruffle
cell leading edge
extrinsic component of membrane
clathrin−coated vesicle
retromer complex
presynaptic endocytic zone
clathrin vesicle coat
clathrin coat
midbody
endocytic vesicle membrane
ruffle membrane
clathrin adaptor complex
Count
p.adjust

**Suppl. Fig. 2**

**Suppl. Figure 2. Proteomic characterization of sucrose cushioning from different neuronal subcellular compartments.** (**a**) Western blot of the cleared lysates (input, 3% of the total volume) or cushion samples (50% of the total volume). (**b**) Quantification of the Western blot analysis. While the core Ribosomal Protein RPS11 was significantly enriched through sucrose cushioning both after 30 min and 210 min long centrifugations, the Ribosome Biogenesis Factor Bystin was significantly enriched only after 210 min long centrifugation. Each dot represents a biological replicate. For RPS11, Anova, $p = 3.7e{-}05$; *t*-test between input and 30min cushion, $p = 0.00019$; *t*-test between input and 210min cushion, $p = 0.0097$. For Bystin, Anova, $p = 2.1e{-}05$; *t*-test between input and 30min cushion, $p = 0.0061$; *t*-test between input and 210min cushion, $p = 0.012$. Both the global p-value (Anova test, on the top) and the p-value for each comparison (t-test) between the input and the cushion samples are shown. Abbreviations: small ribosomal subunit (40S), biological replicate (bio.repl.). (**c**) Bar plot of the number of quantified protein groups in the cushion samples across biological replicates of each subcellular compartment. The bars related to the somata+neurites compartment are colored in dark green, the ones for neurites only in light green. (**d**) Hierarchical clustering of biological replicates according to protein levels in cushion samples from somata+neurites or neurites compartments. Cells are color coded according to Pearson correlation coefficients. (**e**) PCA analysis showing similarities across protein levels in cushion samples from somata+neurites (dark green) or neurites only (light green) compartments. The variability explained by each Principal Component (PC) is shown in brackets. (**f**-**g**) Gene Ontology (GO) analysis of Cellular Components terms overrepresented (FDR<0.01) among the differentially regulated proteins between total lysates and cushion samples from either somata+neurites (**f**) or neurites-only (**g**).

**a**

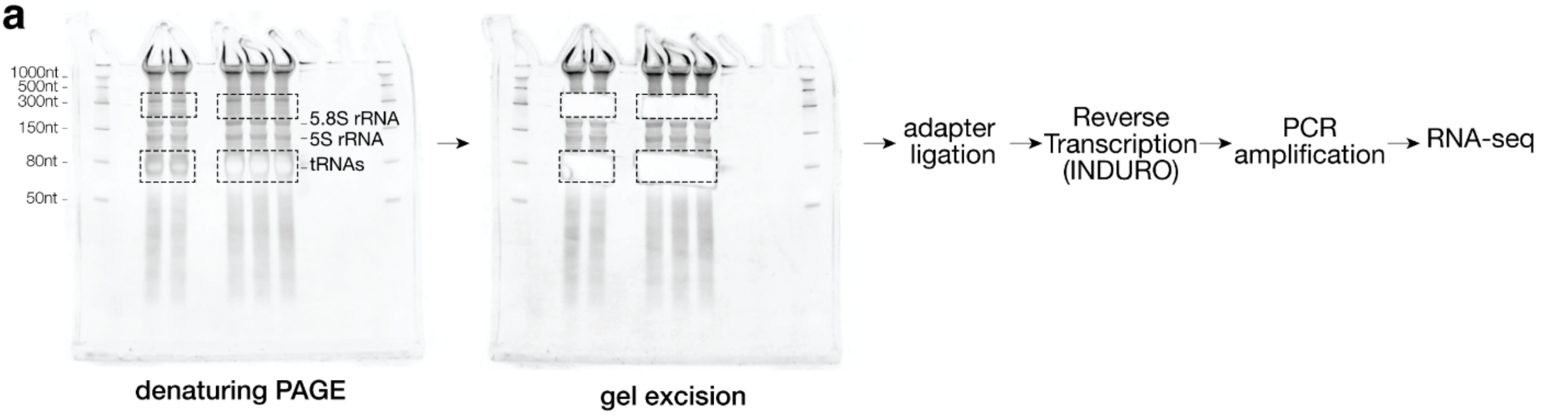


**b**

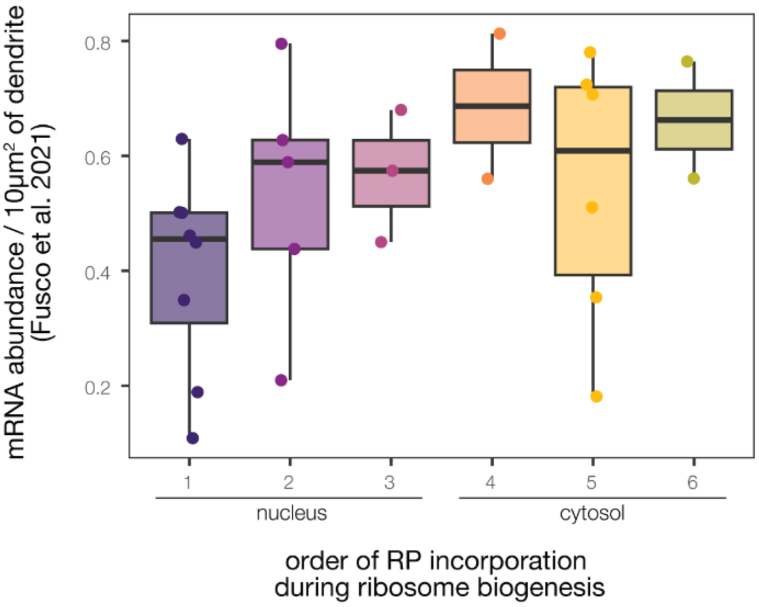


**Suppl. Fig. 3**

**Suppl. Figure 3. Customized pre-rRNA-seq pipeline and RP mRNAs localization in dendrites.** (**a**) Schematic of the pipeline for the pre-rRNA-seq experiments. After running the purified RNA samples in a denaturing gel, RNA species of interest are enriched by gel excision. For the pre-rRNA samples, gels are cut between 160 and 400nt. Samples are then processed for adapter ligation and the Reverse Transcription, both of which introduce a 3'end bias. In particular, the Reverse Transcriptase INDURO is used because it is better able to by-pass secondary structures. After PCR amplification, samples are prepared for RNA sequencing. (**b**) Box plot of the mean RP mRNA abundance per 10μm$^2$ of dendrite (as measured in Fusco et al. 2021) according to the order and site of RP incorporation during ribosome biogenesis. Each dot represents a Ribosomal Protein.
